# Ki-67 contributes to normal cell cycle progression and inactive X heterochromatin in p21 checkpoint-proficient human cells

**DOI:** 10.1101/134767

**Authors:** Xiaoming Sun, Aizhan Bizhanova, Timothy D. Matheson, Jun Yu, Lihua Julie Zhu, Paul D. Kaufman

## Abstract

Ki-67 protein is widely used as a tumor proliferation marker. However, whether Ki-67 affects cell cycle progression has been controversial. Here, we demonstrate that depletion of Ki-67 in human hTERT-RPE1, WI-38, IMR90, hTERT-BJ cell lines and primary fibroblast cells slowed entry into S phase and coordinately downregulated genes related to DNA replication. Some gene expression changes were partially relieved in Ki-67-depleted hTERT-RPE1 cells by co-depletion of the Rb checkpoint protein, but more thorough suppression of the transcriptional and cell cycle defects was observed upon depletion of cell cycle inhibitor p21. Notably, induction of p21 upon depletion of Ki-67 was a consistent hallmark of cell types in which transcription and cell cycle distribution were sensitive to Ki-67; these responses were absent in cells that did not induce p21. Furthermore, upon Ki-67 depletion, a subset of inactive × (Xi) chromosomes in female hTERT-RPE1 cells displayed several features of compromised heterochromatin maintenance, including decreased H3K27me3 and H4K20me1 labeling. These chromatin alterations were limited to Xi chromosomes localized away from the nuclear lamina and were not observed in checkpoint-deficient 293T cells. Altogether, our results indicate that Ki-67 integrates normal S phase progression and Xi heterochromatin maintenance in p21 checkpoint-proficient human cells.

## INTRODUCTION

Ki-67 was first identified via an antibody raised against Hodgkin lymphoma cell nuclei (1). Because Ki-67 is generally expressed strongly in proliferating cells and poorly in quiescent cells (2), anti-Ki-67 antibodies are frequently used to detect proliferative cells in clinical studies (3,4). In interphase cells, Ki-67 primarily localizes to the nucleolus (5-7), whereas during mitosis, it coats the chromosomes (8-10). In the past few years, several studies have greatly increased our understanding of Ki-67 function. This is particularly true for its mitotic roles. Specifically, Ki-67 is required for formation of the mitotic perichromosomal layer (11,12), a proteinaceous sheath that coats mitotic chromosomes (13,14). As part of this layer, Ki-67’s large size and highly positively-charged amino acid composition keeps individual mitotic chromosomes dispersed rather than aggregated upon nuclear envelope disassembly, thereby ensuring normal kinetics of anaphase progression (15). At anaphase onset, Ki-67 binds protein phosphatase 1γ (PP1γ) to form a holoenzyme (16) important for targeting substrates that must be dephosphorylated during mitotic exit (10). In contrast to its structural role on the mitotic chromosomal surface, Ki-67 does not appear to affect nucleosomal spacing (15) or condensation of individual mitotic chromosomes (11,15).

In addition to its expression in proliferating cells, other experiments suggested Ki-67 has a positive role in regulating cell proliferation. In early studies, antisense oligonucleotides targeting Ki-67 expression in human IM-9 multiple myeloma cells blocked [^3^H]-thymidine incorporation, indicative of inhibition of proliferation (17). Likewise, Ki-67-targeted phosphorothioate anti-sense oligonucleotides that resulted in partial depletion of Ki-67 protein inhibited proliferation of human RT-4 bladder carcinoma and other tumor cell lines (18). More recently, siRNA-mediated depletion of Ki-67 resulted in reduced proliferation in human 786-0 renal carcinoma cells (19).

However, despite its utility as a proliferation marker, the contribution of Ki-67 to cell proliferation has recently been questioned. For example, in one recent study, genetic disruption of Ki-67 in human MCF-10A epithelial breast and DLD-1 colon cancer cells did not affect cell proliferation rates in bulk culture, although clonogenic growth of highly diluted cell populations was reduced (20). In another recent study, depletion of Ki-67 in human HeLa or U2OS cells did not alter cell cycle distribution (12). These data raise the possibility that Ki-67 function may have different consequences in different cell types.

In our previous studies we demonstrated that the interphase and mitotic localization of Ki-67 is partially dispersed in cells lacking the N-terminal domain of the p150 subunit of Chromatin Assembly Factor-1 (21, 22). We therefore began exploring the functions of Ki-67 in several human cell types. Here, we show that the contribution of Ki-67 to cell proliferation depends on cell type. In hTERT-RPE1, WI-38, IMR90, and hTERT-BJ cells and primary foreskin fibroblasts, depletion of Ki-67 resulted in reduced frequencies of S-phase cells and concomitant reduction in S phase-related transcript levels. We show that in female hTERT-RPE1 cells, these phenotypes required a p21 checkpoint-mediated delay in S phase entry, and are accompanied by altered nucleolar association and chromatin characteristics of the inactive × (Xi) chromosome. Notably, none of these phenotypes were observed in human cells unable to induce p21 in response to Ki-67 depletion. Therefore, Ki-67 is important for normal S phase progression in p21 checkpoint-proficient human cells, in a manner correlated with its contribution to Xi heterochromatin composition.

## RESULTS

### Ki-67 affects S phase gene expression and progression

To explore how Ki-67 impacts gene expression, we performed RNAseq analyses of control and Ki-67-depleted hTERT-RPE1 cells, a diploid retinal pigment epithelial cell line immortalized by an hTERT transgene (23). Duplicate analyses were highly reproducible (Figures 1A-B). Ki-67 depletion resulted in approximately equal numbers of reduced and increased RNA levels across the transcriptome (Figure 1C). However, Reactome pathway analysis of RNA abundance changes showed that the most altered functional sets of genes included those involved in DNA replication and cell cycle progression (Figure 1D). For example, levels of RNAs encoding all subunits of several protein complexes involved in S phase progression were concertedly reduced, including DNA replication clamp loader RFC (RFC1-5 genes), ssDNA-binding complex RPA (RPA1-3), the replicative helicase (MCM1-6), the GINS replication initiation complex (GINS1-4), the DNA polymerase alpha/primase complex, as well as the DNA replication clamp PCNA and the flap-endonuclease FEN1 involved in Okazaki fragment maturation (Supplemental Tables 1-2). We confirmed reduced levels of a subset of replication-related RNA targets by RT-qPCR analyses (Figures 1E-F). Notably, the RT-qPCR data were very similar when obtained with two distinct Ki-67 depletion reagents, one a synthetic siRNA and the other a cocktail of in vitro-diced dsRNAs non-overlapping the siRNA target, both of which efficiently depleted steady-state Ki67 levels (Figures 1M-N).

**Figure 1.**
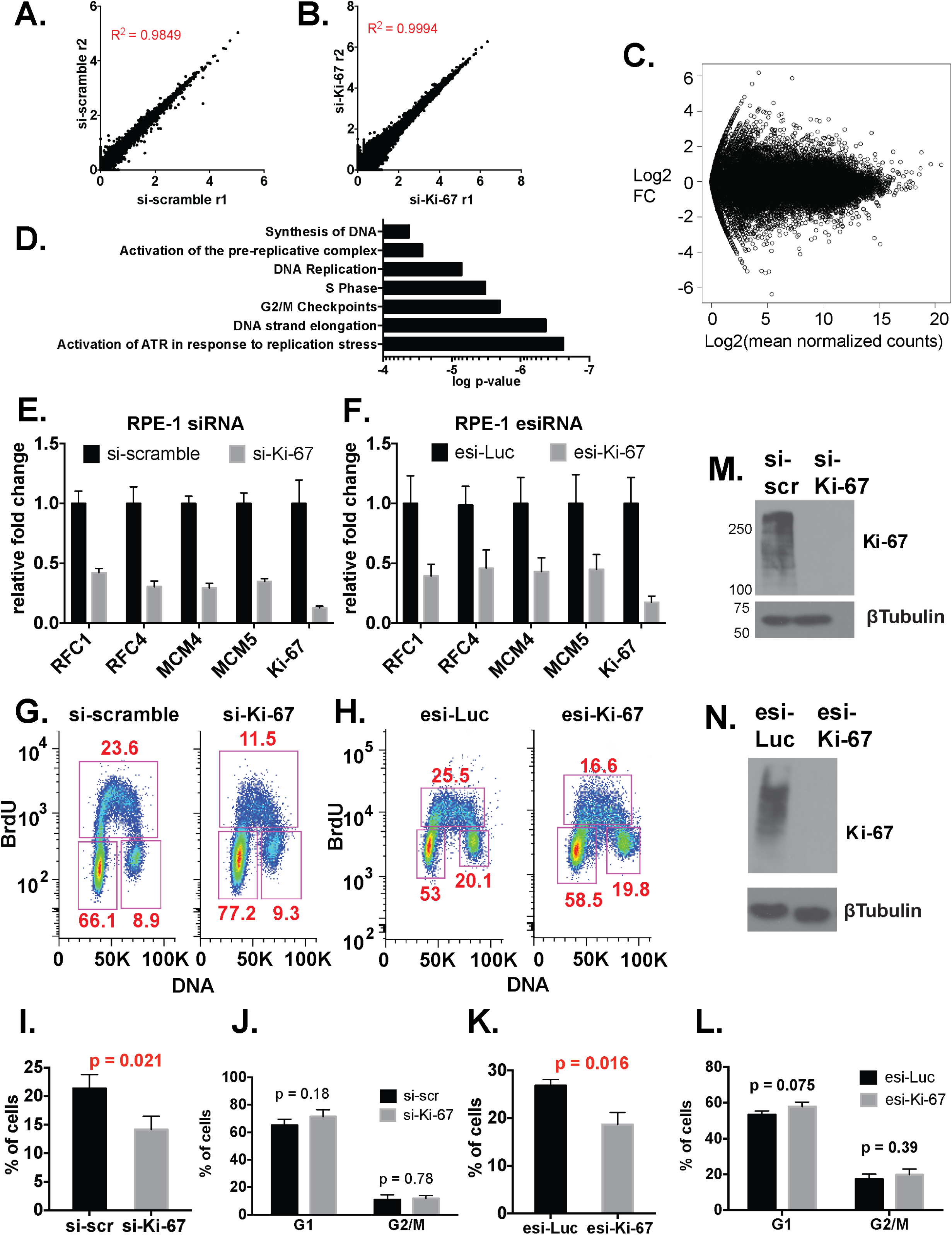
Ki-67 depletion in hTERT-RPE1 cells reduced S-phase-related mRNA abundance and the proportion of cells in S phase. A. Scatter plot analysis of RNA levels, comparing two replicate si-scramble RNAseq analyses of hTERT-RPE1 cells. R: Pearson’s correlation coefficient. B. Scatter plot analysis comparing two replicate siKi-67 RNAseq analyses. C. Distribution of RNA level fold changes (FC) measured by RNAseq, comparing si-scramble and siKi-67-treated hTERT-RPE1 cells. The x-axis shows the mean log2 value for normalized counts of abundance levels for each RNA species. The y-axis shows the log2 fold change upon Ki-67 depletion. The symmetry of the plot above and below the y = 0 axis indicates that similar numbers of genes are up- and down-regulated upon Ki-67 depletion. D. Reactome evaluation of RNAseq analysis of si-Ki-67-treated cells. The PATH terms with a p-value < 5e-05 are graphed. E. RNA levels of DNA replication genes are coordinately down-regulated in siKi-67-treated cells. RT-qPCR measurements are presented as fold change relative to the scramble siRNA control after normalization. *MKI67* mRNA levels indicate effectiveness of the siRNA treatment. Data are mean ± std. dev. of 3 biological replicates. F. Analysis of RNA levels as in panel E, except that cells were treated with in vitro-diced esiRNAs as depletion reagents. G. FACS analysis of siRNA-treated cells. Cells were pulsed with BrdU for 20 min, and analyzed via two-dimensional flow cytometry monitoring BrdU incorporation (y-axis) and DNA content (x-axis). G1 (lower left), G2 (lower right), and S-phase populations (upper box) are boxed in each sample, with percentages of the total population shown. Data shown are from one representative experiment of three biological replicates performed. H. FACS analysis as in panel G, except that cells were treated with esiRNAs. I. Quantification of percentage of cells in S-phase in siRNA-treated hTERT-RPE1 populations from three biological replicate BrdU-labeling experiments. p-value comparing si-scramble and si-Ki-67 treatments is indicated, calculated via an unpaired, two-tailed parametric t test. J. Quantification of percentage of cells in G1 or G2/M phase from the same three experiments analyzed in panel I. K. Quantification of percentage of S-phase cells as in panel I, except that cells were treated with in vitro-diced esiRNAs. L. Quantification of percentage of cells in G1 or G2/M phase from the same three experiments analyzed in panel K. M. Immunoblot analysis of Ki-67 depletion in siRNA-treated hTERT-RPE1 cells from panel D. Marker molecular weights are indicated on the left. N. Immunoblot analysis of Ki-67 depletion in esiRNA-treated hTERT-RPE1 cells from panel F.

We also observed that fluorescence-activated cell sorting (FACS) analysis of Br-dUTP-labeled cells showed that the proportion of S phase cells decreased upon depletion of Ki-67 in hTERT-RPE1 cells with either depletion reagent (Figure 1G-H; quantified in Fig. 1I and 1K). In contrast, proportions of G1 and G2/M populations were not significantly altered (Fig. 1J and 1L). Therefore, Ki-67 is important for normal S phase distribution and gene expression in hTERT-RPE1 cells.

As an additional control for the direct effect of the siRNA treatment, we used CRISPR/Cas9 to mutate the siRNA target site in hTERT-RPE1 cells (Fig. 2A-B). In a resulting homozygous mutant cell line (Figure 2C), siKi-67 no longer depleted Ki-67 protein levels, but as expected the esiKi-67 reagent was still effective (Figure 2D). We also observed that siKi-67 no longer altered candidate S phase RNA levels in the resistant cell line, but esiKi-67 did (Figure 2E). We conclude that the transcriptional response to acute depletion of the Ki-67 mRNA is due to loss of Ki-67 protein.

**Figure 2.**
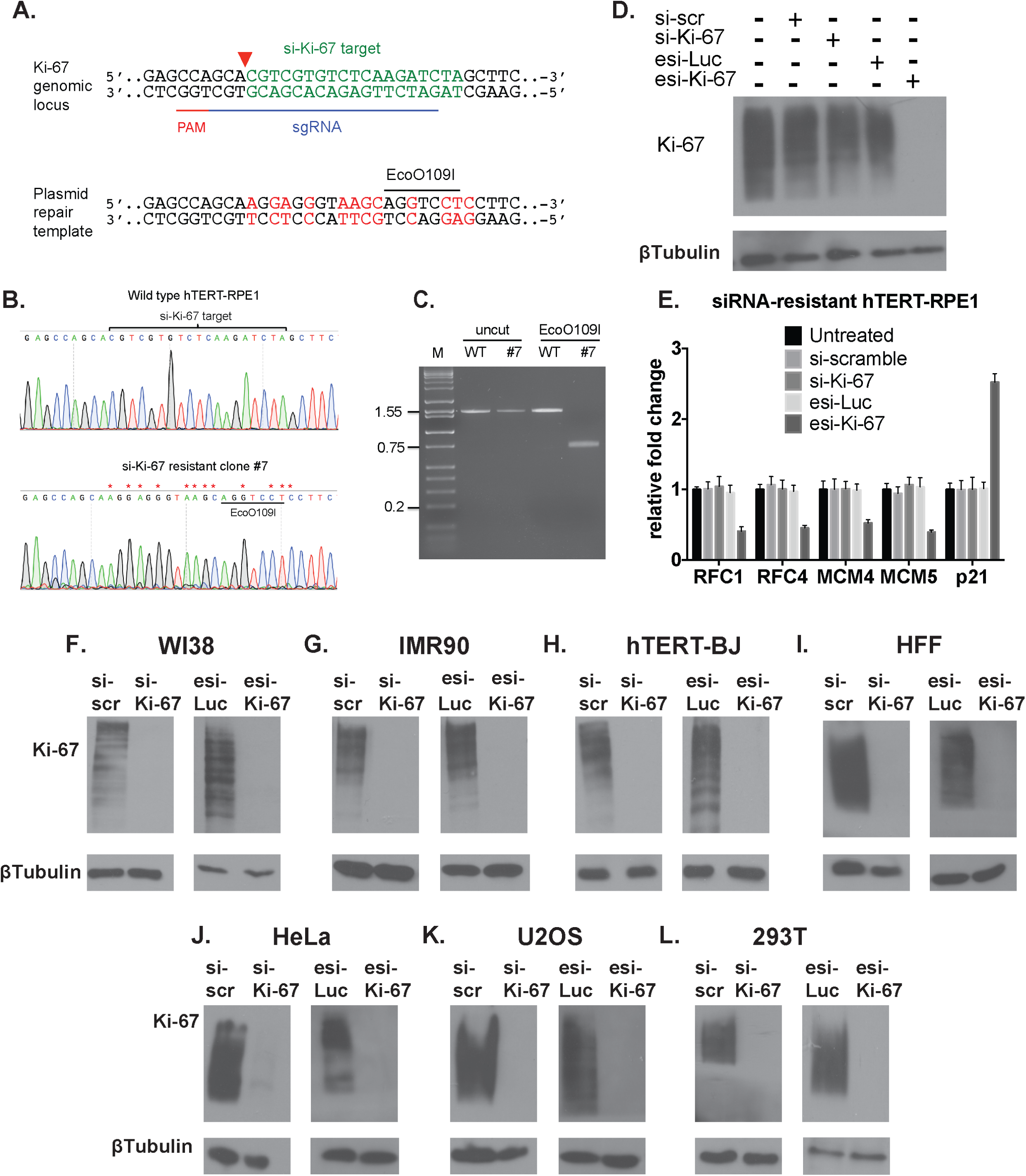
Validation of specificity and effectiveness of Ki-67 depletion reagents. A. CRISPR/Cas9-based strategy for generating siRNA-resistant mutations in the endogenous Ki-67 gene. The si-Ki-67 target (green) and sgRNA-directed cleavage site (red triangle) are indicated on the upper diagram of the endogenous locus. Altered nucleotides (red) and the novel EcoO109I restriction site are shown on the lower diagram of the repair template. B. DNA sequence analysis of a PCR product from wild-type hTERT-RPE1 cells and si-Ki-67-resistant clone #7. C. EcoO109I digestion of the same PCR product sequenced in panel B. D. Immunoblot analysis of clone #7 treated with the indicated reagents. E. RT-qPCR analysis of clone #7. F. F-L. Immunoblot analyses of the indicated cell lines, treated with the indicated RNA depletion reagents.

However, recent studies challenge the view that Ki-67 is important for human cell proliferation; for example, depletion of Ki-67 had minimal effects on the cell cycle distribution of tumor-derived HeLa or U2OS cells (12). Because our data indicated that Ki-67 contributes to normal cell cycle progression in hTERT-RPE1 cells, we hypothesized that the contribution of Ki-67 to cell cycle progression would be cell type-dependent, and may be related to checkpoint function. To explore this idea, we depleted Ki-67 in several additional cell lines. We compared diploid, non-immortal WI-38 and IMR90 fibroblasts, hTERT-immortalized BJ fibroblasts, and primary human foreskin fibroblasts (HFFs) with tumor-derived cell lines: virally-transformed kidney (293T) or cervical carcinoma (HeLa) cells, and osteosarcoma U2OS cells. Importantly, we confirmed that all these cells could be efficiently depleted of Ki-67 protein using both of our depletion reagents (Figure 2F-L).

In all of the experiments with the non-tumor-derived cells, we observed reduced replication factor RNA levels and fewer cells in S phase, similar to our results in hTERT-RPE1 cells, and these results were independent of which Ki-67 depletion reagent was used (Figures 3-4). In contrast, in the tumor-derived cell lines, Ki-67 depletion did not result in uniform down regulation of the S phase genes tested, nor were there changes in cell cycle distribution (Figures 5-6). These data are consistent with RNAseq data sets in HeLa and U2OS cells (12) that did not display concerted downregulation of DNA replication genes. We conclude that the effects of Ki-67 depletion on RNA levels and cell cycle distribution are cell type-dependent.

**Figure 3.**
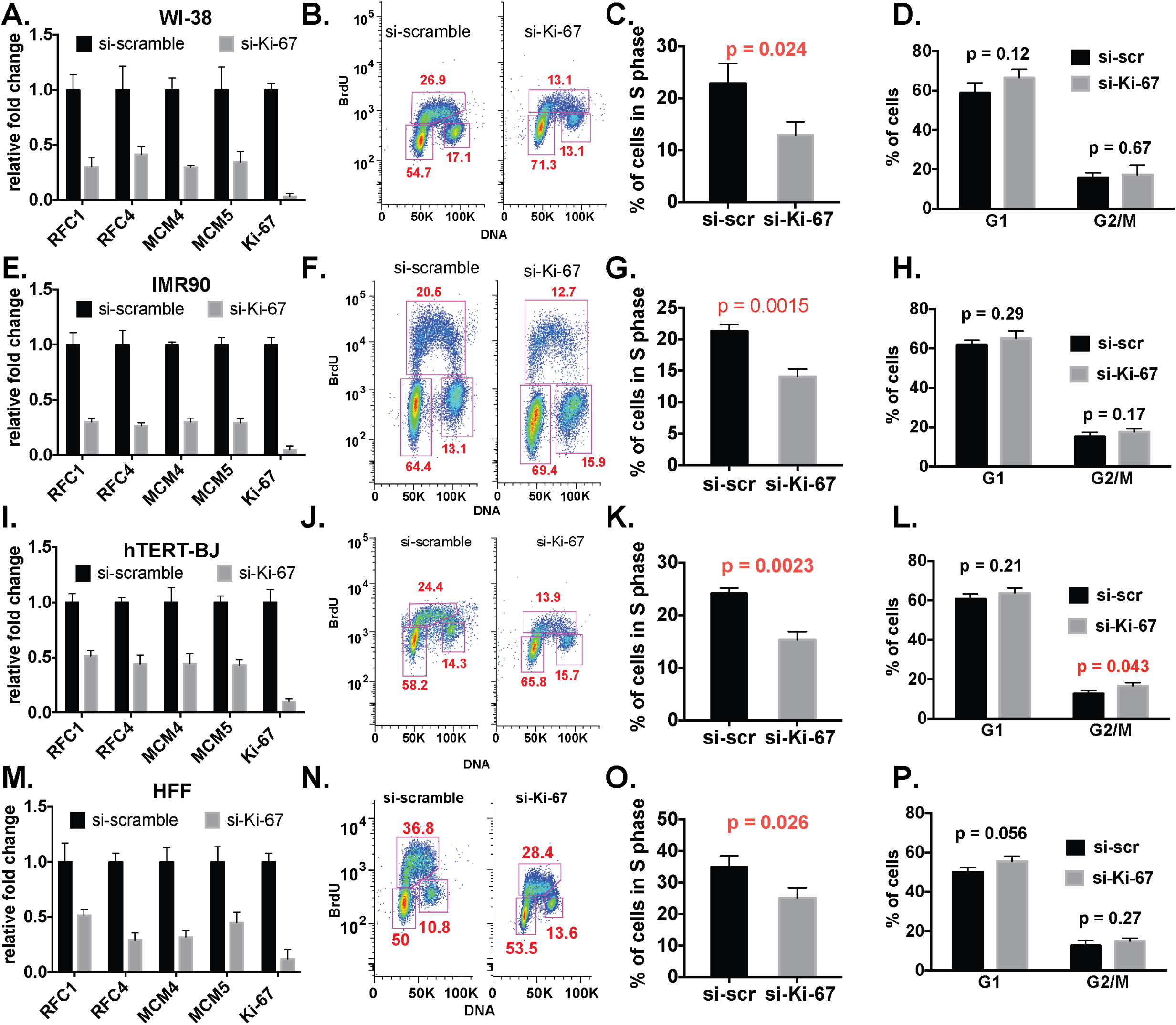
siRNA-mediated Ki-67 depletion affected S-phase gene expression and cell cycle distribution in diploid human cells. WI-38 (panels A-D), IMR-90 (panels E-H), hTERT-BJ (panels I-L) cell lines and human primary fibroblasts (HFF, panels M-P) were analyzed. A, E, I, M. RT-qPCR analysis, as in Figure 1E. B, F, J, N. FACS analysis, as in Figure 1G. C, G, K, O. Quantification of percentage of S-phase cells, as in Figure 1I. D, H, L, P. Quantification of percentage of G1 and G2/M phase cells as in Figure 1J.

**Figure 4.**
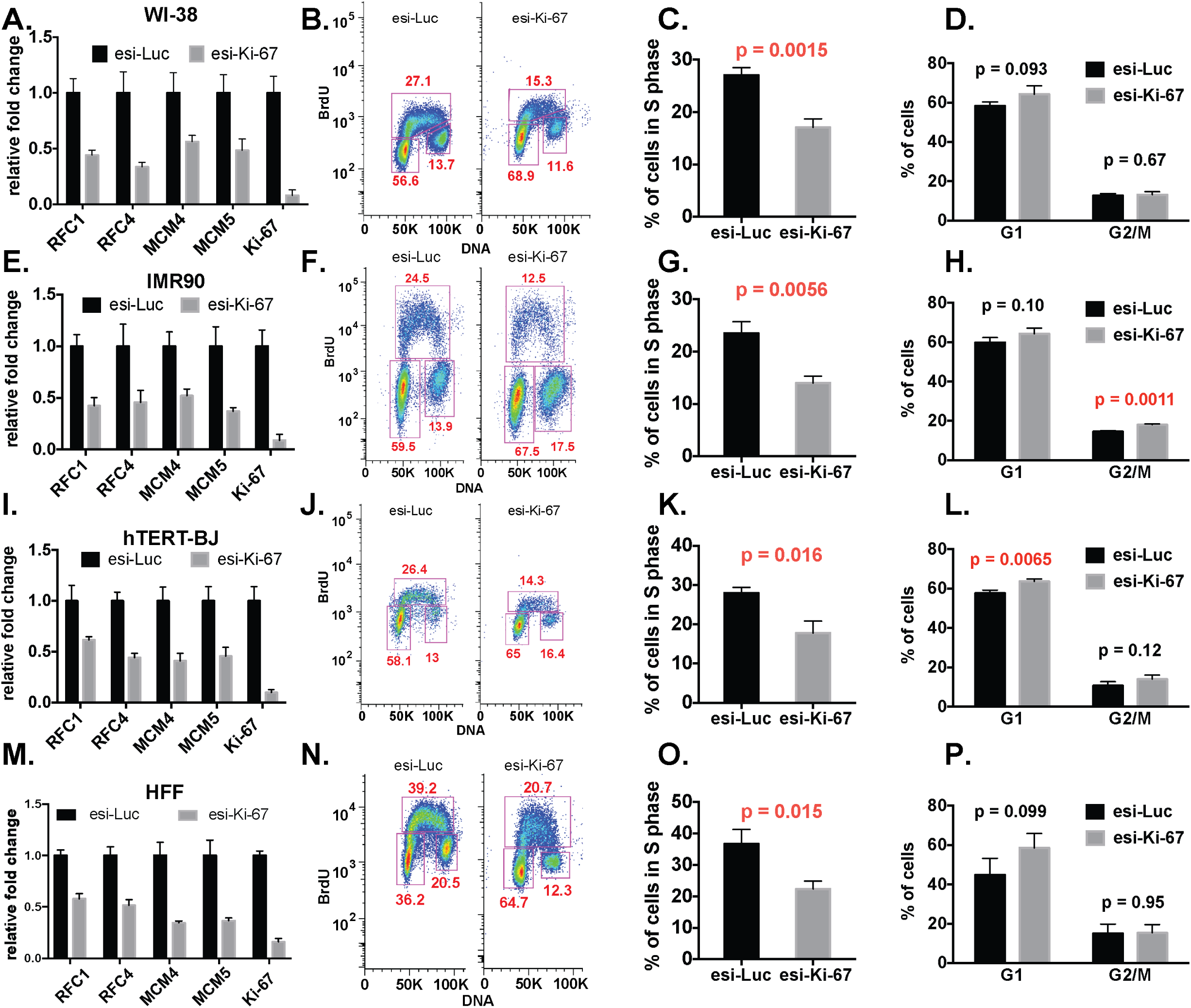
esiRNA-mediated depletion of Ki-67 in diploid cells resulted in the same phenotypes observed with siRNA treatments. Cells and assays were the same as in Figure 3.

**Figure 5.**
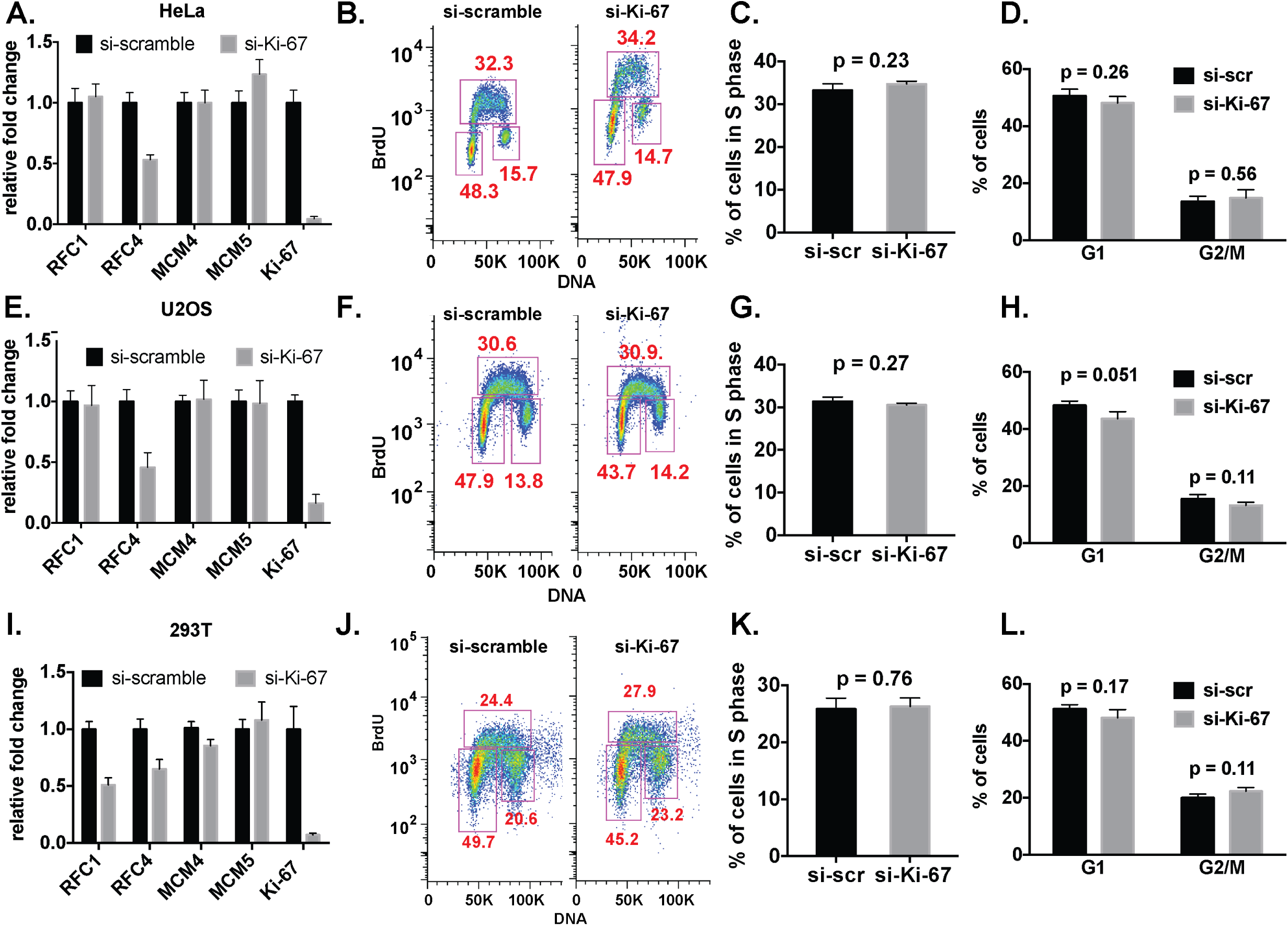
Ki-67-insensitive cells. Ki-67 depletion did not affect S-phase gene expression and cell cycle distribution in HeLa (panels A-D), U2OS (panels E-H), and 293T (panels I-L) cell lines. Panels A, E, I: RT-qPCR analysis. Panels B, F, J: FACS analysis. Panels C, G, K: Quantification of percentage of S-phase cells. Panels D, H, L: Quantification of percentage of G1 and G2/M phase cells.

**Figure 6.**
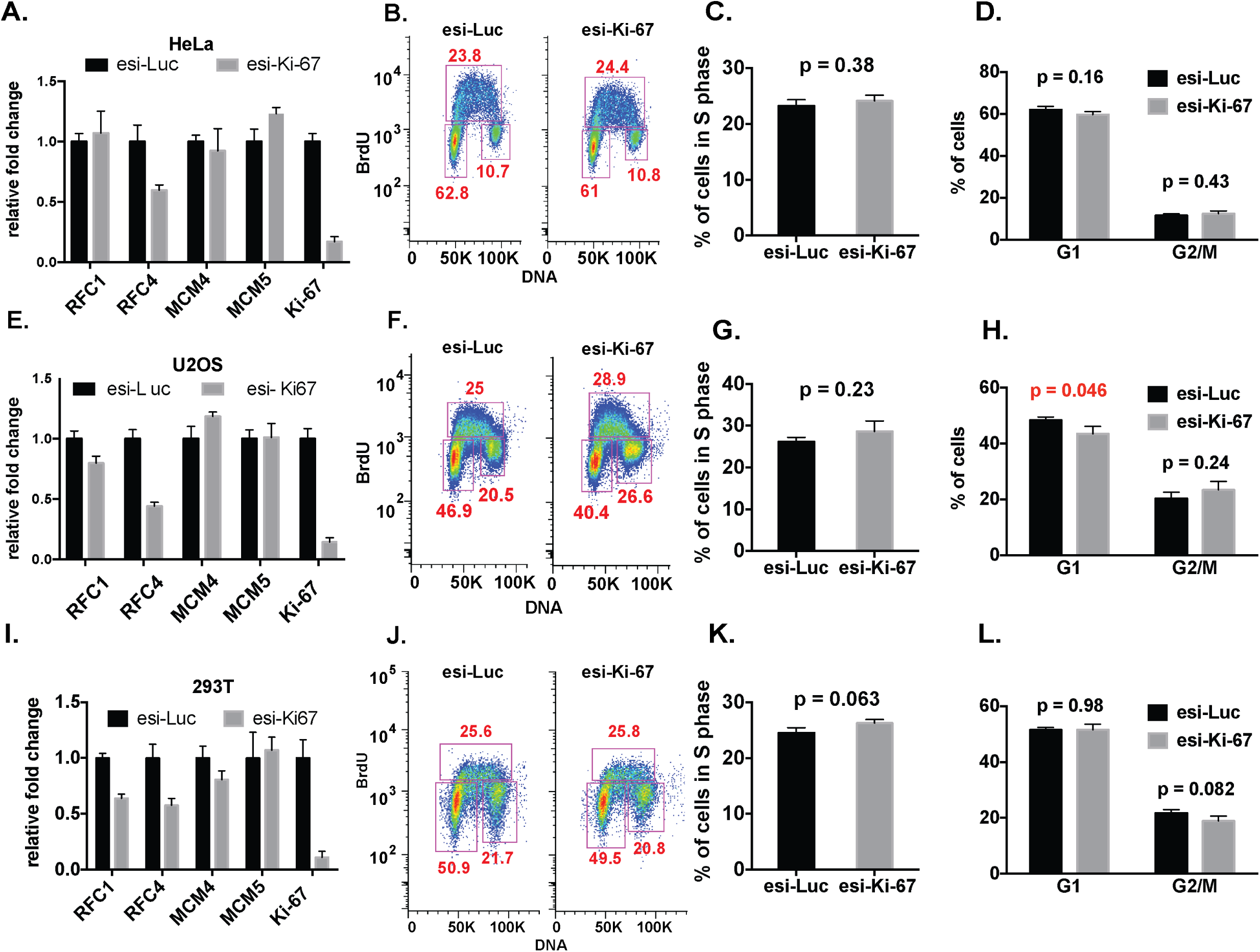
esiRNA-mediated depletion of Ki-67 in insensitive-cells resulted in the same phenotypes observed with siRNA treatments. Cells and assays were the same as in Figure 5.

The FACS analyses of asynchronous cell populations indicated that Ki-67 depletion most significantly affects S phase in the sensitive cell types. To examine this in more detail, we analyzed the kinetics of DNA synthesis in synchronized hTERT-RPE1 cells. Cells were blocked near the G1-S transition of the cell cycle with hydroxyurea (HU) for 15 hours (Figure 7A) (24), which provided efficient arrest (Fig. 7D). Cells were then released into drug-free media and pulse-labeled with the deoxynucleotide analog EdU at two-hour timepoints across an 8-10-hour time course. In control cells treated with the scrambled siRNA, EdU labeling was first detected at the 2 hour time point, and displayed a typical early S phase pattern consisting of many small foci (25). At later time points, the pulse of EdU labeled larger foci, indicative of mid-late S phase patterns. Upon Ki-67 depletion, we observed a delay in the initial detection of EdU incorporation of approximately 2 hours (Figure 7B). Notably, the Ki-67-depleted population also displayed a higher percentage of cells that did not incorporate EdU during the time course (Figure 7C). Together, these data are consistent with our transcriptomic and FACS data indicating that Ki-67 depletion affects S-phase in hTERT-RPE-1 cells (Figure 1).

**Figure 7.**
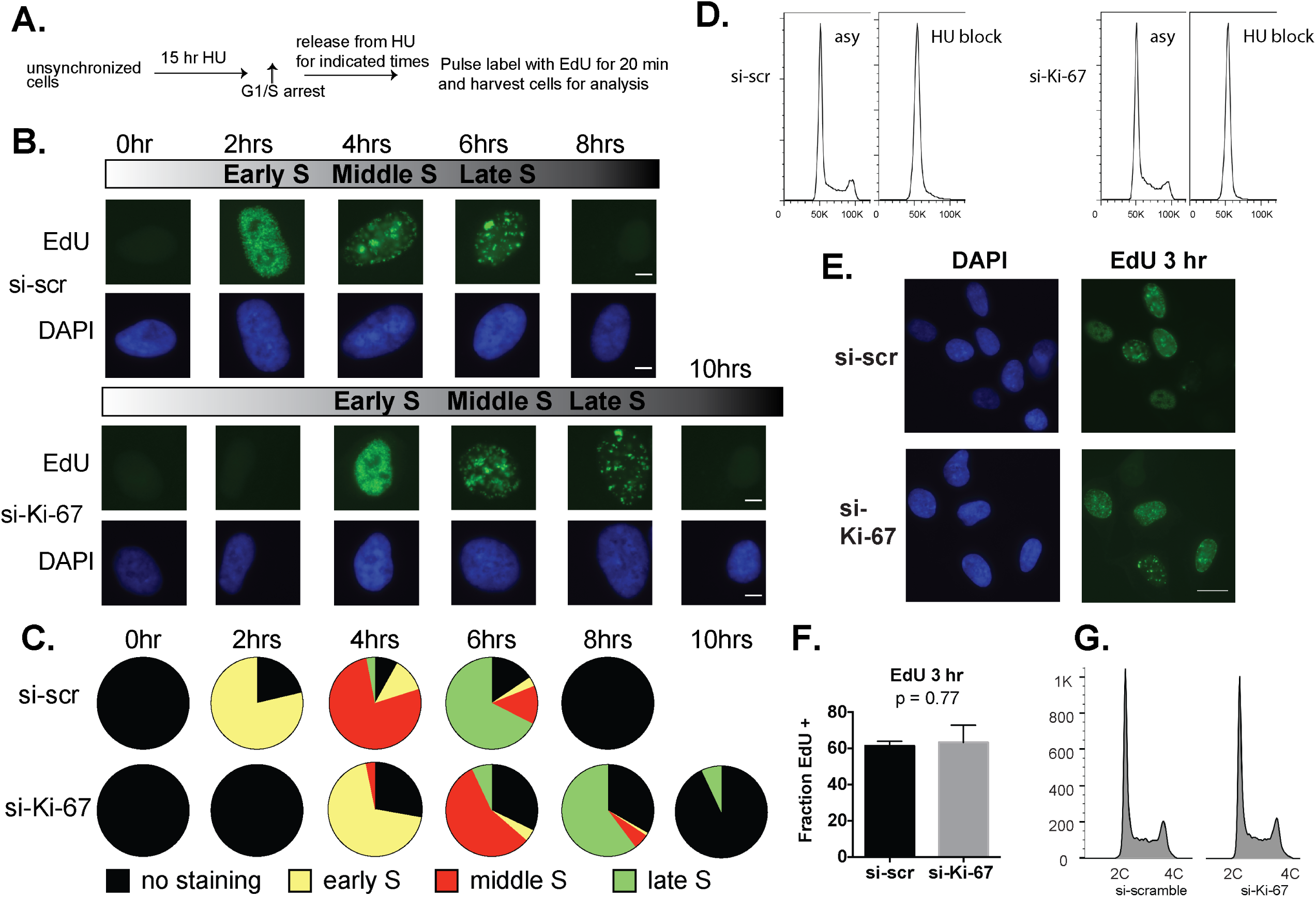
Depletion of Ki-67 delayed S-phase entry in hTERT-RPE1 cells. A. Schematic of short-pulse assay. hTERT-RPE1 cells were released from HU arrest for the indicated time periods, pulsed for 20 min with EdU, and analyzed by click chemistry and fluorescence microscopy for EdU incorporation. B. Representative cells from the indicated time points, showing EdU staining (green) to detect the progression of S phase. Total DNA was visualized with DAPI staining (blue). S phase cells were categorized into 3 sub-stages based on the number, size, shape and distribution of fluorescent-labeled replication foci. During early S phase, small and numerous replication foci were scattered in the nuclear interior, but excluded from nucleolus, nuclear periphery and other heterochromatic regions. At mid-S phase, replication takes place at the nuclear periphery and perinucleolar regions. Late in S phase, there are several large foci throughout the nucleus (25). Scale bar: 5 μm. C. Distributions of EdU morphologies during the time course. Percentages of early S phase, middle S phase and late S phase EdU staining morphologies were counted in > 300 total cells per time point. D. FACS histograms showing cell-cycle profiles of propidium-iodide stained hTERT-RPE1 cells. Asy: Asynchronous cells were incubated with DMSO vehicle control. HU block: cells were incubated with hydroxyurea. Left panel: si-scramble-treated cells were incubated with or without 2mM hydroxyurea for 15 hours. Right panel: siKi-67-treated cells were incubated with or without 2mM hydroxyurea for 15 hours. Histograms were generated using FlowJo v9.9.4. E. A longer (three hour) EdU pulse prevents detection of S phase alterations in Ki-67-depleted hTERT-RPE1 cells. Asynchronous cells were treated with either si-scramble or siKi-67 for 72 hrs, incubated with 5-ethynyl-2-deoxyuridine (EdU) for the final 3 hours, and analyzed via click chemistry. The total cells assayed = 218 for si-scramble and 265 for siKi-67, in three independent experiments. Scale bar, 20 μm F. Ratio of EdU-positive cells to the total cell numbers. ns: not significant (p-value = 0.77). G. Cell cycle distribution of the si-scramble and siKi-67 treated hTERT-RPE1 cells as analyzed by one-dimensional FACS profiling of propidium iodide-stained cells.

### Checkpoint responses to Ki-67 depletion

Because Ki-67 depletion did not affect S phase transcription or cell cycle progression in tumor-derived cell lines, our data suggested that functional checkpoints are required for sensitivity to Ki-67 depletion. Consistent with this idea were comparisons of our RNAseq data with metadata analyses of genes regulated by cell cycle status or by E2F transcription factors (26) that are important for G1/S cell cycle phase transcription (26-28). These meta-analyses aggregated multiple datasets, finding that similar results in multiple datasets strongly predicted regulatory network connections that could be missed in single experiments. Of the cell cycle-regulated genes identified in that study, we find those that peak during G1/S phase were more frequently downregulated than upregulated upon Ki-67 depletion (Fig. 8A; Supplemental Table 3). Consistent with this observation, E2F target RNA levels (Fig. 8B) were much more frequently downregulated than upregulated upon Ki-67 depletion. These comparisons were consistent with the idea that checkpoint activation contributed to the observed delay in S phase entry and transcriptional phenotypes in Ki-67-depleted cells.

**Figure 8.**
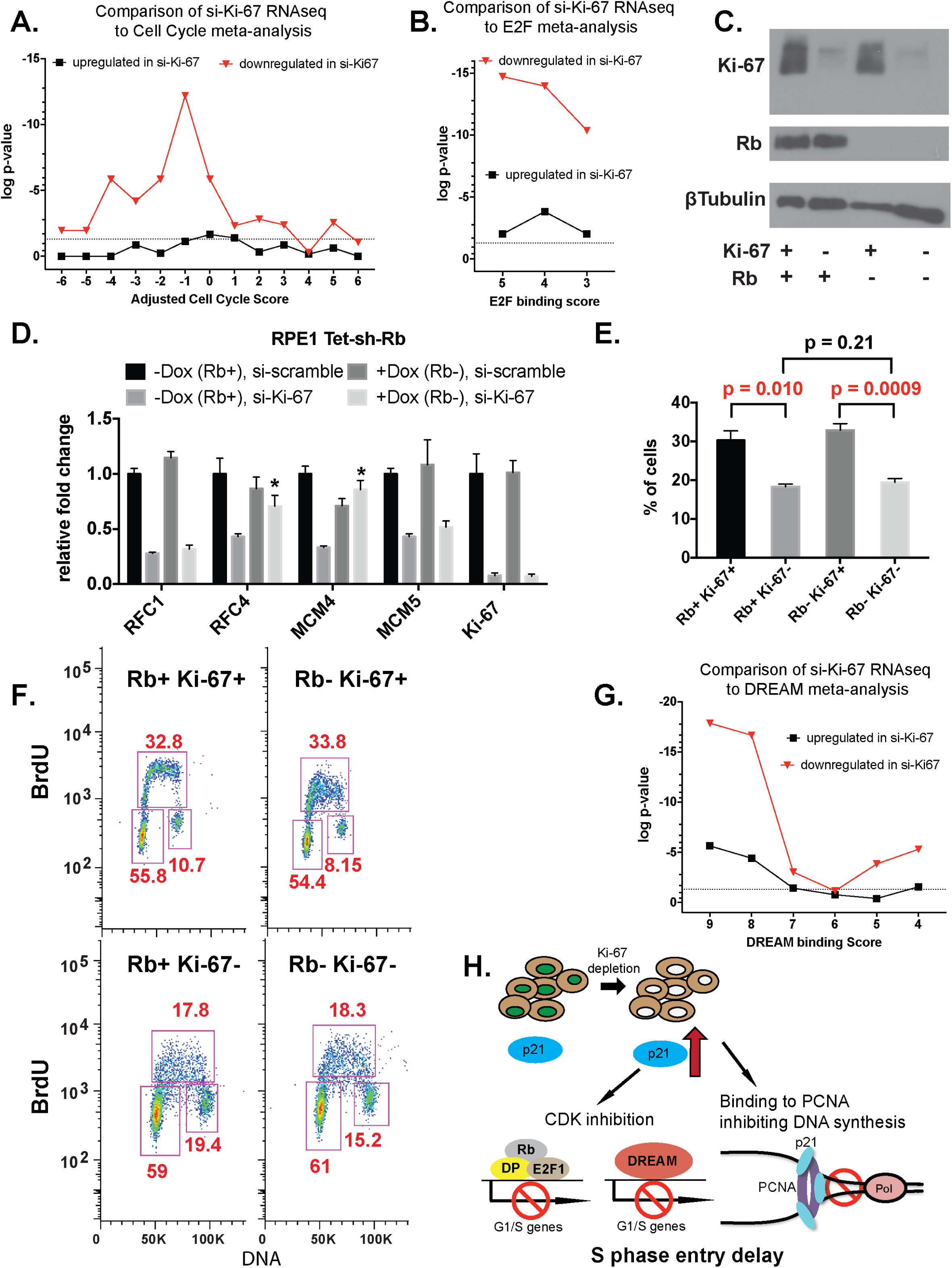
Rb contributes to transcriptional downregulation caused by Ki67 depletion. A. Summary of transcriptional changes of cell cycle target genes (based on Table S10 in (26)). The “Adjusted Cell Cycle Score” on the y-axis indicates values based on meta-analysis of 5 different cell cycle expression data sets, plus information regarding binding by Rb/E2F and MMB/FOXM1 transcription factors. Negative values indicate frequent detection of G1/S expression and binding by Rb/E2F, and positive values indicate frequent detection of G2/M expression and binding by MMB-FOXM1. B. p-values of transcription changes of E2F target genes (based on Table S9 in (26)), with the greater score on the x-axis representing higher frequency of detection as an E2F target. As expected from panel A, E2F targets are commonly downregulated upon Ki-67 depletion. C. Immunoblot analysis hTERT-RPE1 Tet-sh-Rb cells. Cells were treated with either vehicle (Rb+) or 2 μg/ml doxycycline (Rb-) as indicated for 72 h to induce sh-Rb expression, and were also incubated in the presence of either si-scramble (Ki-67+) or siKi-67 (Ki-67-). D. RT-qPCR analysis of DNA replication genes in cells treated as in panel C. Measurements are presented as fold change relative to the scramble siRNA control without doxycycline induction. Data are mean ± std. dev. of 3 biological replicates. p-values were calculated via unpaired, two-tailed parametric t tests and corrected for multiple comparisons using the Holm-Sidak method. E. Percentage of S phase cells calculated from three BrdU labeling experiments. p-values were calculated via unpaired, two-tailed parametric t tests. F. FACS analysis of a representative BrdU labeling experiment. G. p-values of transcription changes of DREAM target genes (based on Table S7 in (26)), with the greater score on the x-axis representing more frequent detection as a DREAM target. H. Model. In “Ki-67 sensitive” cells, depletion of Ki67 leads to p21 induction. The elevated p21 levels are predicted to downregulate the Rb/E2F and DREAM target G1/S cell cycle genes and inhibit DNA synthesis by binding to PCNA. Together, these effects delay S phase entry.

To test this, we performed experiments co-depleting checkpoint proteins. First, we took advantage of a derivative of hTERT-RPE1 cells that have an integrated, doxycycline-inducible shRNA that targets the RB mRNA (29). Rb levels remained unchanged in these cells in the absence of doxycycline (Fig. 8C), and siRNA-mediated depletion of Ki-67 resulted in reduced S phase-related RNA levels as was observed previously (Fig. 8D). Addition of doxycycline to deplete Rb, together with a control scrambled siRNA leaving Ki-67 levels unchanged, did not significantly alter S-phase related target RNA levels (Fig. 8D). In contrast, simultaneous depletion of Rb and Ki-67 resulted in RNA levels at two of the four loci tested that were significantly elevated compared to those in cells depleted of Ki-67 alone (Fig. 8D). FACS analysis indicated that Rb depletion was insufficient to significantly change the cell cycle profile of Ki-67-depleted cells (Fig. 8E-F). We conclude that depletion of Rb only partially relieves the cellular response to Ki-67 depletion in hTERT-RPE1 cells.

Therefore, we reasoned other factors must contribute. One clue was provided by comparison of the si-Ki-67 RNAseq data to metadata analysis of binding by subunits of the transcription repressor complex termed DREAM (26). We observed that genes with the highest predicted probability of DREAM binding were very frequently downregulated upon Ki-67 depletion (Fig. 8G). In mammalian cells, DREAM is a master regulator of cell cycle-dependent gene expression, repressing both G1/S and G2/M targets, and gene repression by DREAM requires the p21 checkpoint protein (27,30,31). p21 is a potent universal CDK inhibitor (CKI). During G1 and S phases, p21 directly binds to and inhibits the kinase activity of cyclin E-CDK2, cyclin B1-CDK1 and cyclin D-CDK4/6 complexes (32-34). Furthermore, p21 also directly inhibits DNA synthesis by binding to PCNA, the sliding clamp required for processive DNA polymerase activity (35). Therefore we hypothesized that p21 could be important for the response to Ki-67 depletion (Figure 8H).

Consequently, we next tested effects of Ki-67 depletion the *CDKN1A* gene, which encodes p21. Our RNAseq data indicated increased *CDKN1A* RNA levels in Ki-67 depleted hTERT-RPE1 cells (log2 fold change = +0.48, p = 0.016), although multiple hypothesis testing indicated that these values did not achieve the stringent statistical significance cutoff of q < 0.05 (Supplemental Table 2). RT-PCR measurements of *CDKN1A* RNA levels demonstrated a significant increase in four diploid cell types, but not in 293T cells (Fig. 9A). We note that induction of *CDKN1A* RNA was dependent on having an intact siRNA target site in the Ki-67 gene, indicating this is a direct effect of Ki-67 depletion (Figure 2E). Consistent with the increased RNA levels, we detected elevated p21 protein levels in hTERT-RPE1 but not 293T cells (Fig. 9B).

**Figure 9.**
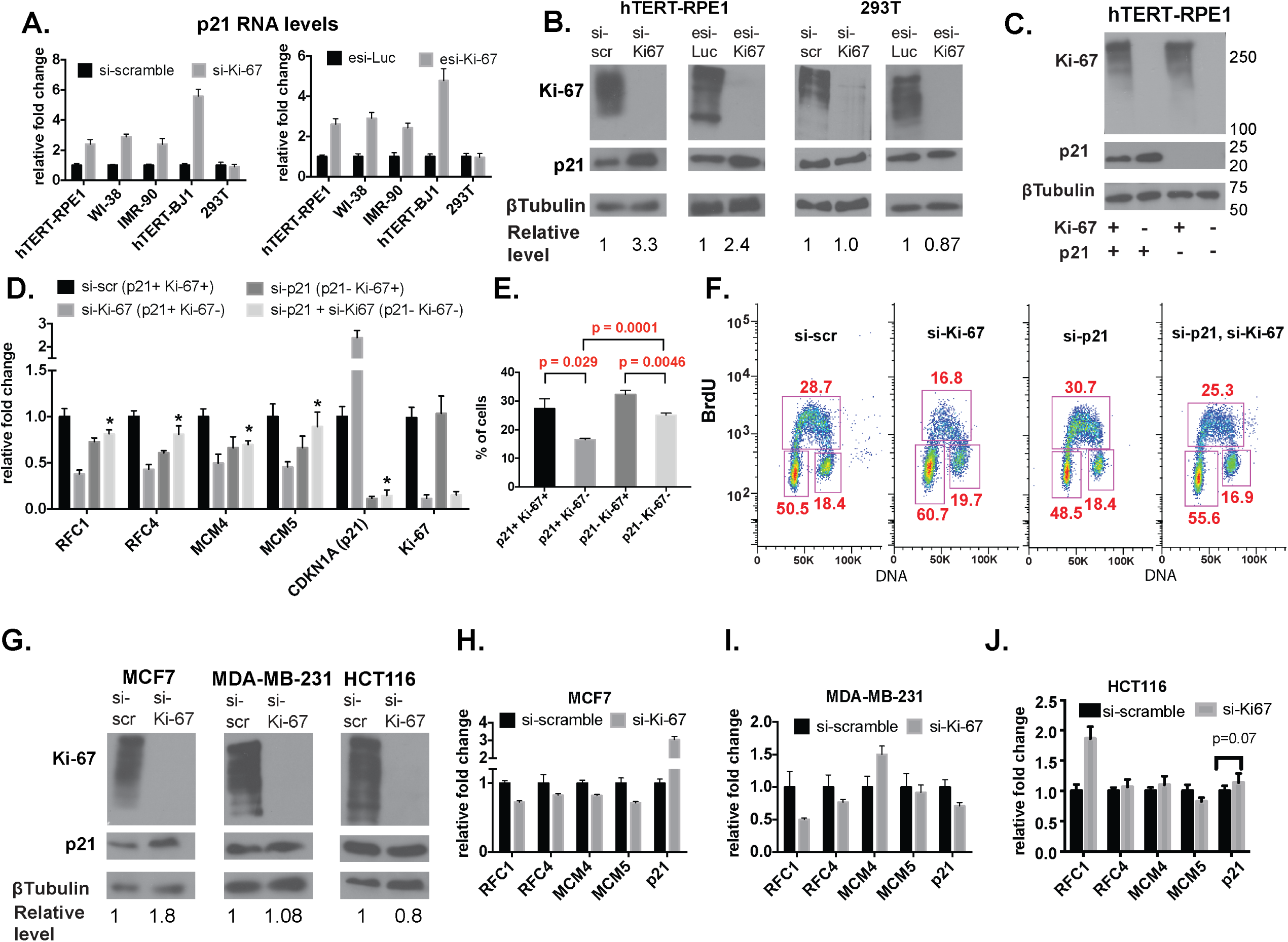
Cell-type-specific induction of a p21-dependent checkpoint upon depletion of Ki-67. A. RT-qPCR analysis demonstrates cell-type specific induction of *CDKN1A* (p21) RNA upon Ki-67 depletion. (left panel) Indicated cells were treated with either si-scramble or si-Ki-67 for 72 hrs. (right panel) Same, except cells were treated with esiRNAs. B. Cell-type specific induction of p21 protein levels upon Ki-67 depletion. hTERT-RPE1 and 293T cells were treated with the indicated siRNA or in vitro-diced esiRNA depletion reagents. p21 protein levels were quantified as a ratio to ß-tubulin and normalized to levels in control-treated cells. Quantification was done in Image J 10.2. C. Immunoblot analysis of hTERT-RPE1 cells depleted of the indicated proteins via siRNA treatments. Marker molecular weights are indicated on the right. D. RT-qPCR analysis of DNA replication genes in hTERT-RPE1 cells from panel C. Asterisks indicate values in the si-p21 + si-Ki-67 samples that were significantly different (p < 0.05) from the siKi-67 samples. p-values were calculated via unpaired, two-tailed parametric t tests and corrected for multiple comparisons using the Holm-Sidak method. E. Percentage of S phase cells calculated from three BrdU labeling experiments. p-values were calculated via unpaired, two-tailed parametric t tests. F. FACS analysis of a representative BrdU labeling experiment. G. Immunoblot analysis of additional siRNA-treated cell lines testing for p21 induction. H. RT-qPCR analysis of the indicated genes in MCF7 cells. p-values calculated as in panel D were < 0.05 for all genes. I. RT-qPCR analysis of the indicated genes in MDA-MB-231 cells. J. RT-qPCR analysis of the indicated genes in HCT-116 cells.

p21 is a direct target of transcriptional induction by the tumor suppressor p53, and the cell lines examined thus far therefore implicated active p53 in the sensitivity to Ki-67 (36,37). To examine this relationship further, we compared the effects of Ki-67 depletion in additional cancer cell lines, including two expressing wild-type p53, MCF7 and HCT116, and also MDA-MB-231 cells which express a mutant p53 defective for p21 induction (38). In MCF7 breast adenocarcinoma cells, we observed that Ki-67 depletion elevated p21 RNA and protein levels (Fig. 9 G,H), and down-regulated replication-related RNAs (Fig. 9H). In contrast, in HCT116 colorectal carcinoma and MDA-MB-231 breast adenocarcinoma cells, Ki-67 depletion did not increase p21 expression or cause concerted down-regulation of S phase genes (Fig. 9G, I, J). Because HCT116 and MDA_MB-231 cells differ in their p53 status, we conclude that p53 status cannot always predict the response to Ki-67 depletion. Instead, we find that induction of p21 upon Ki-67 depletion is a consistent hallmark of this form of checkpoint activation.

To assess the functional consequence of p21 induction, we performed co-depletion experiments in hTERT-RPE1 cells (Fig. 9D). We observed that cells simultaneously depleted of both Ki-67 and p21 no longer displayed the reduced levels of any of the four S phase-related mRNAs analyzed (Fig. 9D). Furthermore, FACS analysis of the co-depleted cells showed that there was significant restoration of the percentage of cells in S phase upon codepletion of p21 with Ki-67 (Fig. 9E-F). We conclude that induction of p21 is functionally important for the effects of Ki-67 depletion on cell cycle distribution in hTERT-RPE1 cells.

### Ki-67 affects heterochromatic characteristics of the inactive X chromosome

Ki-67 is required for the normal cellular localization of heterochromatin-associated histone modifications (12), and for the interphase nucleolar association of heterochromatic loci(22). Because the inactive X (Xi) chromosome is a well-studied example of facultative heterochromatin that associates with the nucleoli of female mouse (24) and human cells (39-41), we tested whether Ki-67 affected characteristics of Xi heterochromatin. Indeed, Ki-67 depletion in hTERT-RPE1 cells resulted in a subset of cells that displayed reduced staining intensity for antibodies recognizing H3K27me3 and H4K20me1, histone modifications that are enriched on the Xi (42,43)(Figure 10A, C, E, G). H3K27me3 is generated by the Polycomb PRC2 complex and is a keystone of facultative heterochromatic silencing (44-46). H4K20me1 is generated by the PR-Set7/Set8/KMT5a enzyme (47) and together with H3K27me3 is an early mark on Xi chromosomes during the process of XIST-mediated inactivation (43,47). Notably, changes to either of these histone modifications were only observed in cells in which the Xi was localized away from the nuclear periphery (Fig. 10B, D, F, H). Furthermore, these changes were not observed in 293T cells that also lacked the cell cycle response to Ki-67 depletion (Fig. 11). Therefore, the response of hTERT-RPE1 cells to Ki-67 depletion involves two classes of correlated events that are both absent in 293T cells: checkpoint-mediated perturbation of S phase, and altered Xi heterochromatin.

**Figure 10.**
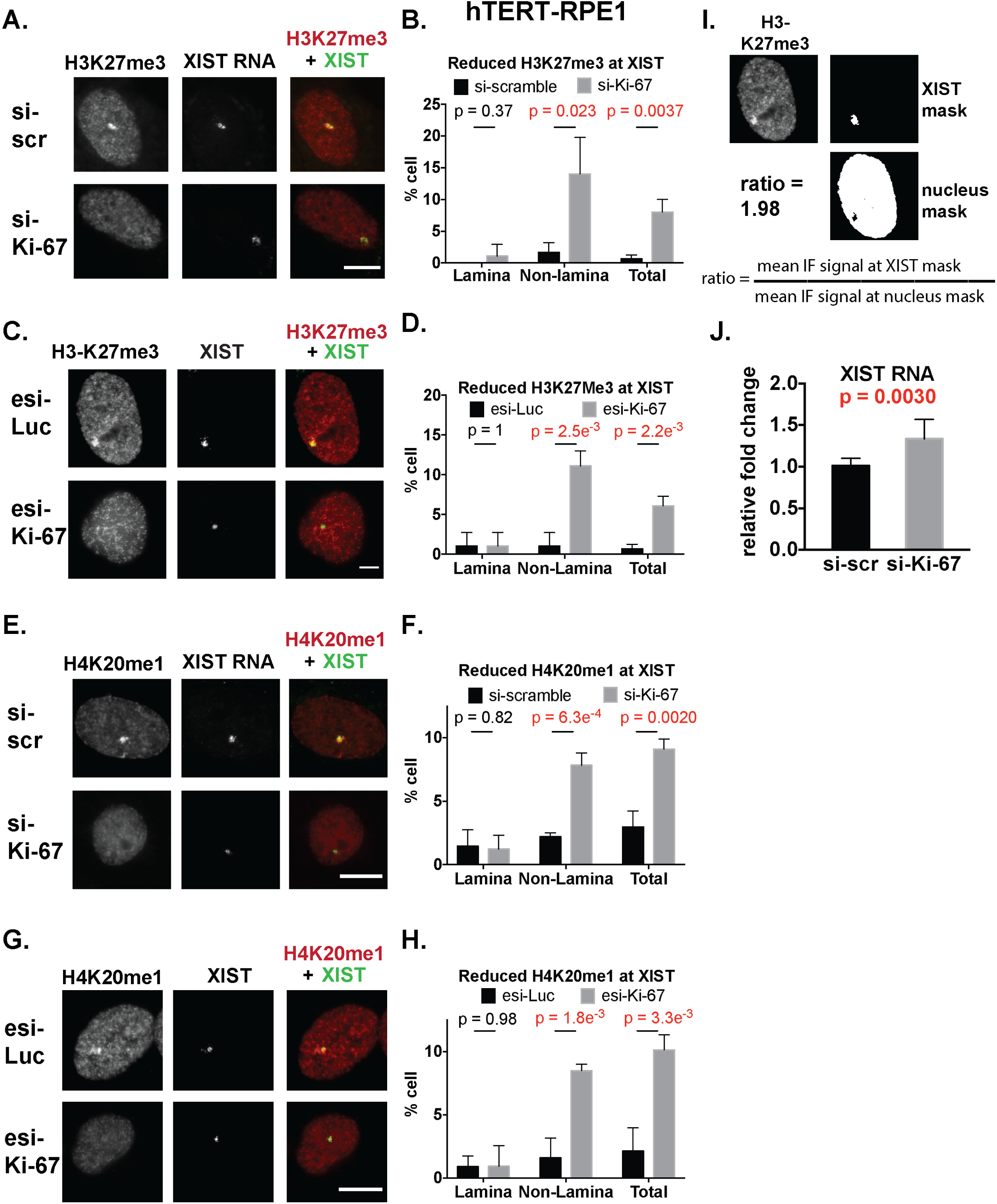
H3K27me3 and H4K20me1 staining of the inactive X chromosome was altered upon Ki-67 depletion in a subset of hTERT-RPE1 cells. Scale bars, 5 μm. A. Immuno-FISH analysis of H3K27me3 overlap with XIST in siRNA-treated hTERT-RPE1 cells. Note that in the si-Ki-67-treated cell, the H3K27me3 signal overlapping XIST displayed reduced intensity and was localized away from the nuclear lamina. B. Quantitation of percentage of cells that display reduced H3K27me3 enrichment on the Xi in the panel A experiments. Enrichment was calculated as the ratio of the mean H3K27me3 signal overlapping XIST divided by the mean H3K27me3 signal from remainder of the entire nucleus. Cells with ratios less than 1.5 were defined as having reduced enrichment, as described previously (48). Percentages were calculated for the total cell populations, as well as for the nuclear lamina-associated XIST foci and the non-lamina-associated foci, as indicated. Total cells assayed = 250 for si-scramble and 239 for si-Ki-67. Mean and SDs are graphed from three biological replicate experiments. p-values were determined by unpaired student’s t tests. C. Analysis of H3K27me3 enrichment on Xi as in panel A, for hTERT-RPE1 cells treated with in vitro-diced esiRNAs. D. Quantitation as in panel B. Total cells assayed= 236 for esi-luciferase, 220 for esi-Ki-67. E. Immuno-FISH analysis of H4K20me1 overlap with XIST in siRNA-treated hTERT-RPE1 cells. Note that the H4K20me1 signal colocalizing with XIST is reduced in the Ki-67-depleted cell. F. Quantitation of the panel E experiments, as in panel B. The total cells assayed = 204 si-scramble and 216 for si-Ki-67. G. Analysis of H4K20me1 in cells treated with esiRNAs. H. Quantitation of the panel G experiments. Total cells assayed = 164 for esi-luciferase, 182 for esi-Ki-67. I. Example of H3K27me3 signal intensity quantification. J. RT-qPCR analysis of XIST RNA levels in hTERT-RPE1 cells.

**Figure 11.**
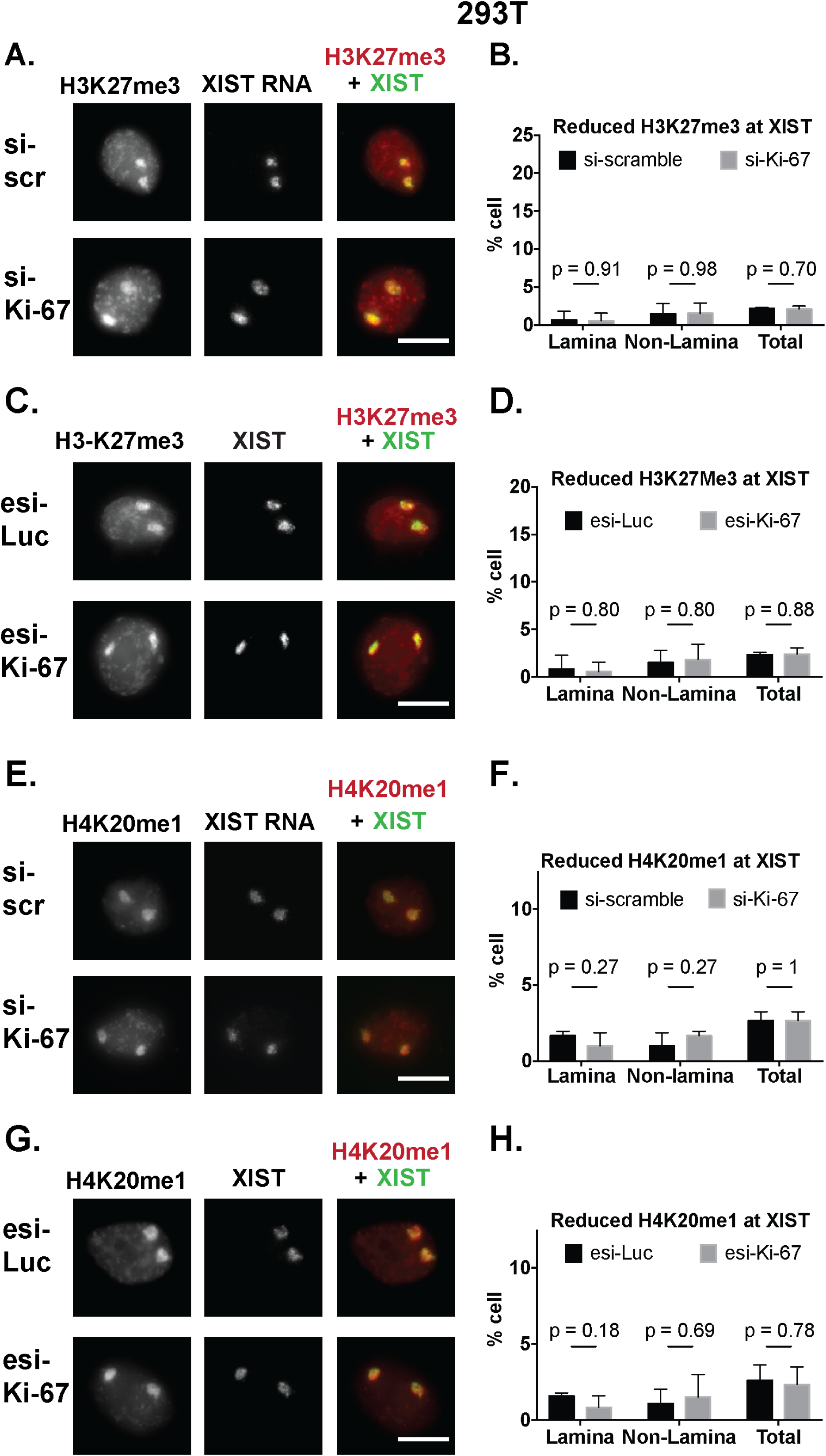
H3K27me3 and H4K20me1 staining of the inactive X chromosome was unaltered upon Ki-67 depletion in 293T cells. Scale bars, 5 μm. A. Immuno-FISH analysis of H3K27me3 overlap with XIST in siRNA-treated 293T cells. Note that 293T cells have two Xi chromosomes. In these cells, the H3K27me3 foci overlapping XIST remained unchanged upon Ki67 depletion. B. Quantitation of panel A. The total alleles assayed = 136 for si-scramble and 146 for si-Ki-67. C, D. Analysis of H3K27me3 in esiRNA-treated 293T cells. Total alleles assayed = 196 for esi-luciferase, 198 for esi-Ki-67. E. Immuno-FISH analysis of H4K20me1 overlap with XIST in 293T cells. In these cells, the H4K20me1 foci overlapping XIST remained unchanged upon Ki67 depletion. F. Quantitation of panel E. The total alleles assayed = 180 for si-scramble and 162 for si-Ki-67. G, H. Analysis of H4K20me1 in esiRNA-treated cells. Total alleles assayed = 180 for esi-luciferase, 162 for esi-Ki-67.

Increased levels of repetitive element-rich Cot-1-hybridizing transcripts and RNA polymerase II have previously been observed in breast cancer cell lines that display perturbations in Xi chromatin (48). We tested for changes in these properties as well, and we detected an increase in the frequency of cells that display Cot-1-hybridizing transcripts or RNA polymerase II on the Xi upon Ki-67 depletion in hTERT-RPE1 cells (Figures 12-13). As observed above for the histone modifications (Fig. 10), increased levels of Cot-1 RNA and Pol II within the XIST domain were only observed in cells in which the Xi was localized away from the nuclear periphery. Also, as for all other phenotypes detected, these changes were similar with either Ki-67 depletion reagents (Figs. 12-13).

**Figure 12.**
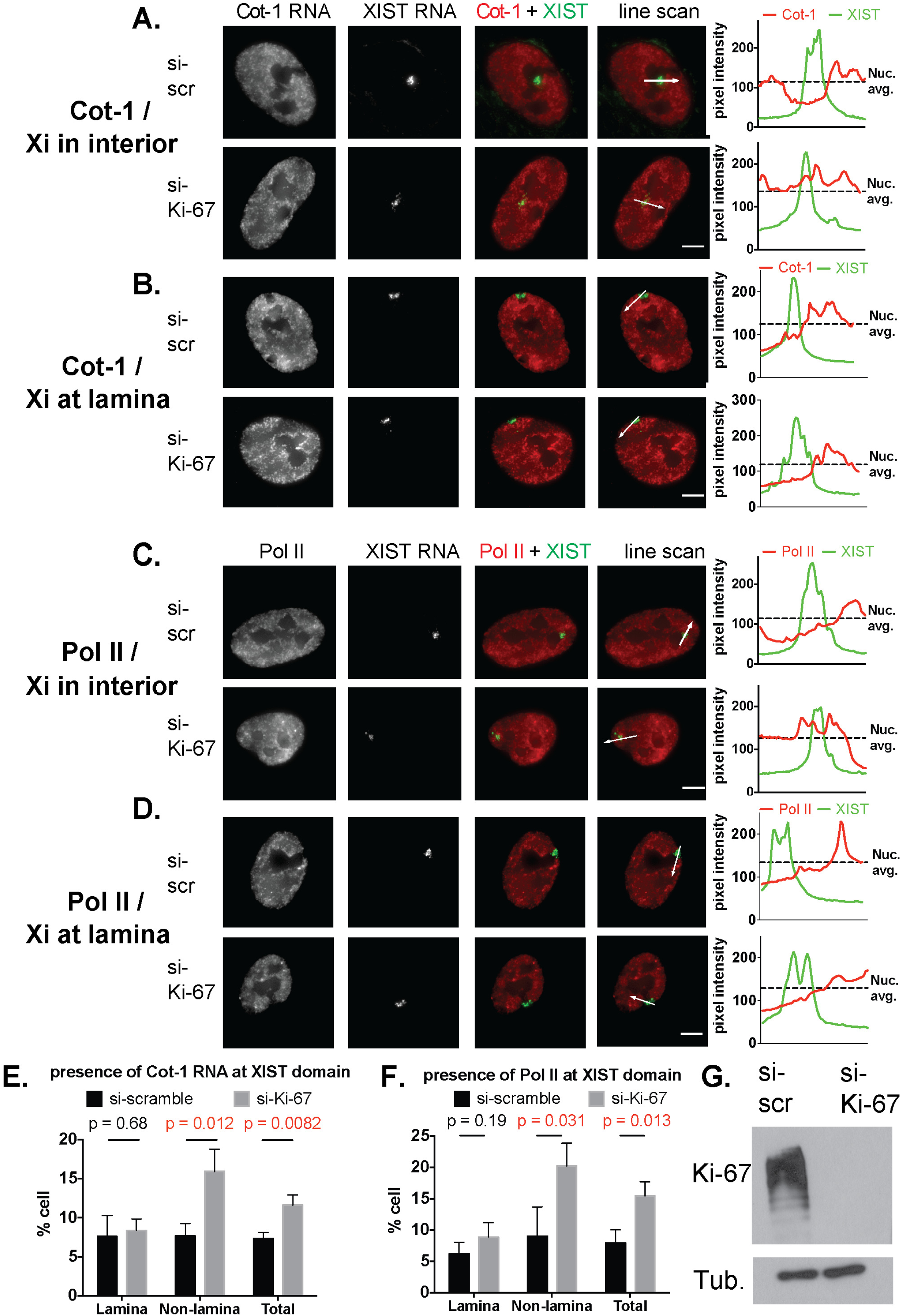
Analysis of Cot-1 and Pol II enrichment on Xi in siRNA-treated hTERT-RPE1 cells. Scale bars, 5 μm. A-B Localization of Cot-1-hybridizing transcripts relative to XIST domains, when XIST is localized away from (A) or at (B) the nuclear lamina. A line scan (white arrow) across the XIST signal (green) was used to analyze Cot-1 hybridization levels (red); fluorescent densities across the line scan were plotted in the right-hand panels. Cot-1 RNA was considered to be reduced across the XIST domain when the average Cot-1 signal overlapping XIST was lower than the average Cot-1 signal across the nucleus. The average nucleus Cot-1 signal is depicted in dotted line. In the examples shown where Xi was within the cell interior (panel A), Cot-1 RNA was excluded from XIST in the si-scramble-treated cell, but not the siKi-67-treated cell. In contrast, siKi-67 treatment did not affect Cot-1 enrichment on Xi in cells where Xi was at the lamina (panel B). C-D Analysis of RNA Pol II localization (red) relative to XIST (green) when XIST is localized away from (panel C) or at (panel D) the nuclear lamina. Exclusion was analyzed as in panels A-B. E. Quantitation of Cot-1 RNA overlap with XIST RNA domains. Mean (and std. dev.) percent of cells displaying Cot-1 RNA overlapping XIST foci are plotted from three biological replicate experiments. The total cells assayed = 404 for si-scramble and 465 for si-Ki-67. p-values were determined by unpaired student’s t tests. F. Quantitation of percentage of cells showing presence of RNA Pol II at Xist RNA domain. The total cells assayed = 362 for si-scramble and 367 for si-Ki-67. G. Immunoblot analysis of Ki67 depletion in siRNA-treated hTERT-RPE1 cells from above.

**Figure 13.**
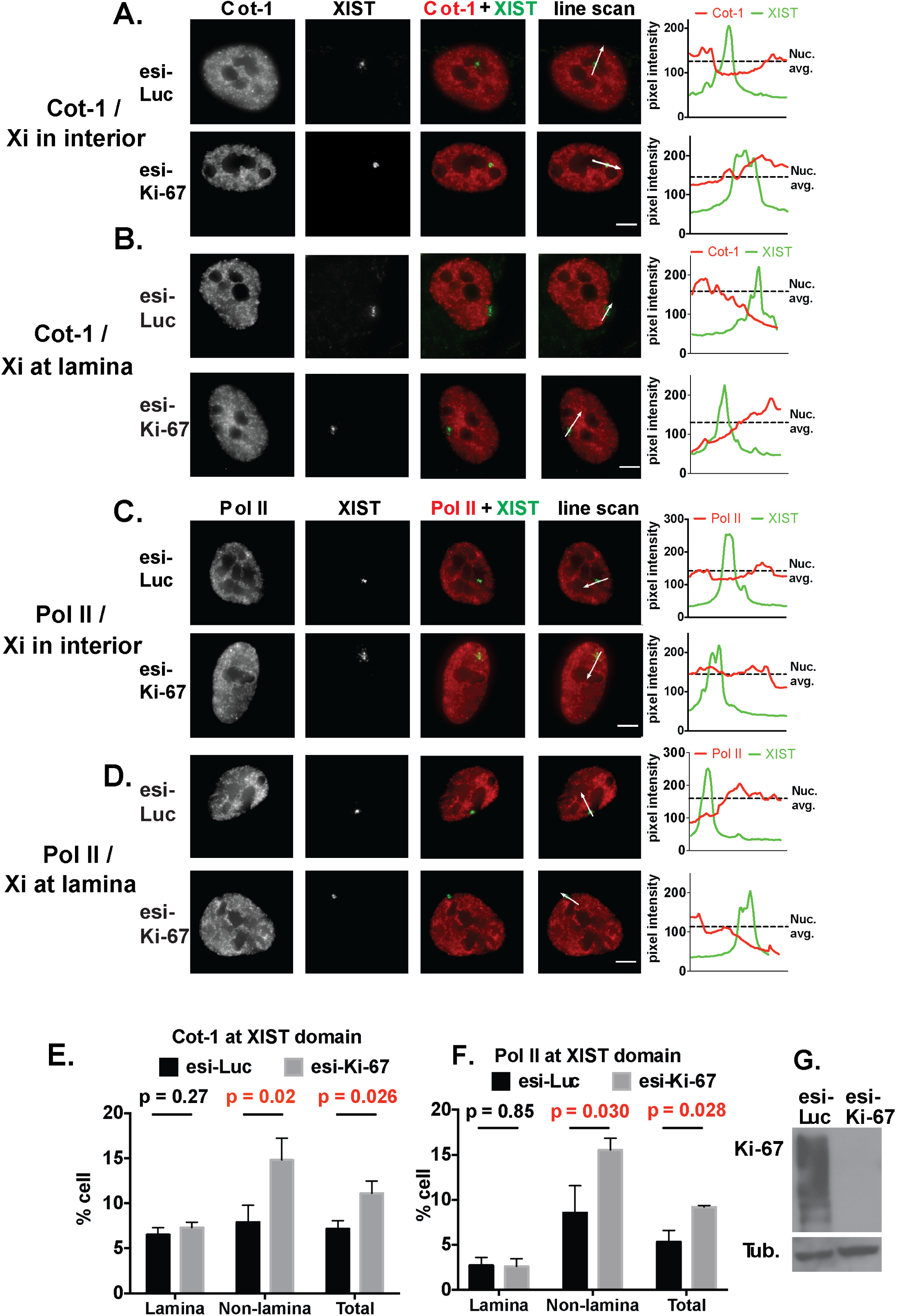
Analysis of Cot-1 and Pol II enrichment on Xi in esiRNA-treated hTERT-RPE1 cells. Analyses were performed as in Figure 12. Scale bars, 5 μm. A, B. Cot-1. Total cells assayed= 178 for esi-luciferase, 180 for esi-Ki-67. C, D. Pol II. Total cells assayed= 160 for esi-luciferase, 176 for esi-Ki-67. E. Quantitation of Cot-1 RNA overlap with XIST RNA domains from panels A-B. F. Quantitation of RNA Pol II at Xist RNA domain from panels C-D. G. Immunoblot analysis of Ki-67 depletion in esiRNA-treated hTERT-RPE1 cells from above.

However, not all aspects of Xi heterochromatin were sensitive to Ki-67 depletion. For example, Ki-67 depletion did not alter the overall appearance of the XIST “cloud” that covers the Xi (Fig. 14A). In addition, RT-PCR showed no significant down regulation of XIST transcript expression (Fig. 10J). Also, there was no evidence for Xi chromosome-wide deprepression of transcription, as shown in the analysis of X-linked gene expression (Fig. 14B), or in the analysis of allele-specific transcription of X-linked genes detected via analysis of known SNPs (Supplemental Table 4). Furthermore, an additional mark associated with the Xi, the histone variant macroH2A, did not change in appearance upon depletion of Ki-67 (Fig. 14C-H). Together, the Xi data indicate that acute depletion of Ki-67 alters several, but not all, characteristics of Xi heterochromatin in hTERT-RPE1 cells. Importantly, changes in H3K27me3 and H4K20me1 staining were not observed in 293T cells, indicating a correlation between checkpoint activation and effects on the Xi upon Ki-67 depletion.

**Figure 14.**
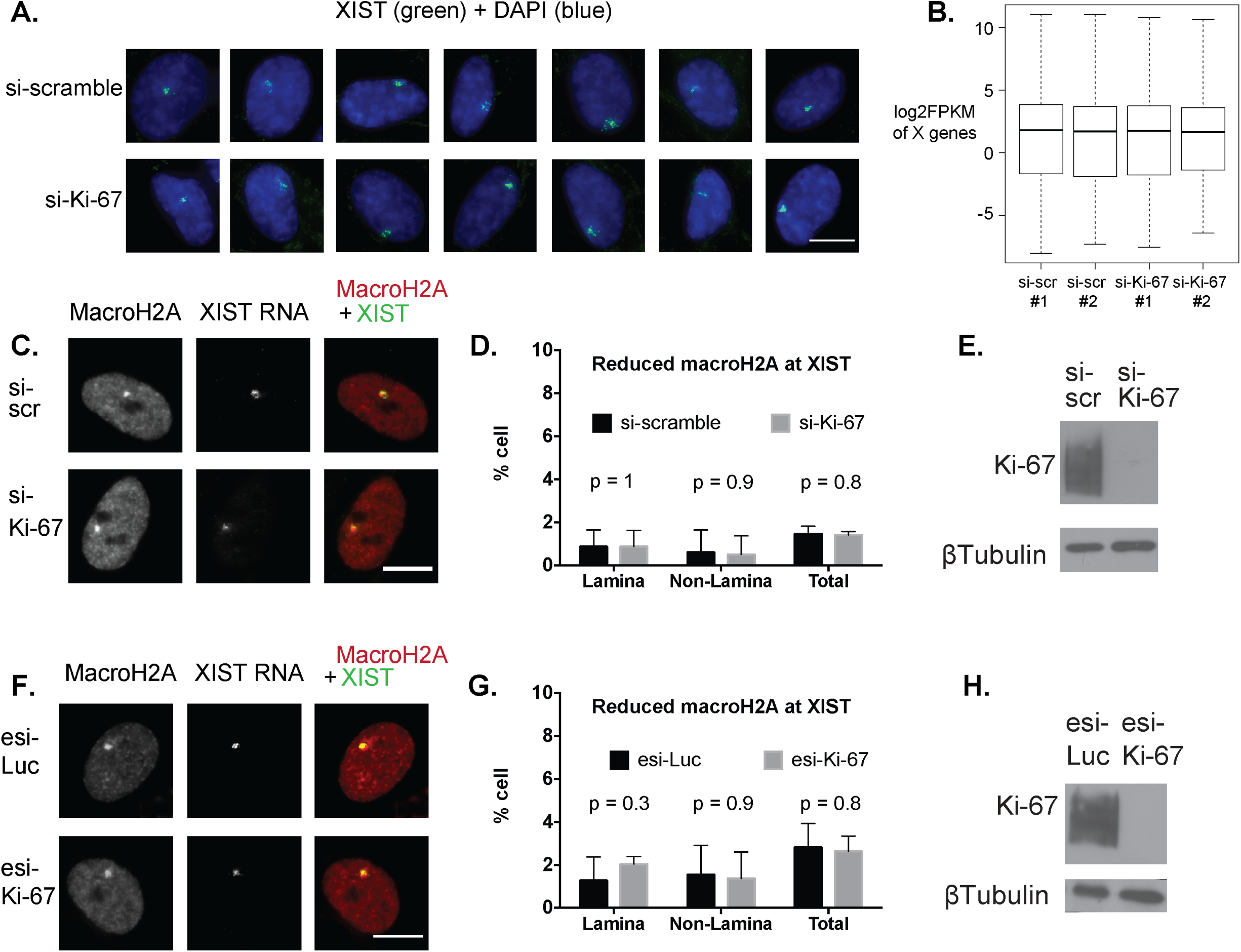
Some aspects of Xi structure and function were resistant to Ki-67 depletion. A. XIST cloud in hTERT-RPE1 cells has similar appearance regardless of Ki-67 depletion. Cells were treated with the indicated siRNAs for 72 hrs and analyzed by RNA-FISH to localize XIST (green) and DAPI staining (blue). Scale bar: 10 μm. B. Average RNA levels of × linked genes did not change upon Ki-67 depletion in hTERT-RPE1 cells. log2 FPKM analyses of RNA-seq data from two biological replicates for the two indicated siRNA treatments are shown. C-H. MacroH2A enrichment at the XIST domain was not altered upon Ki-67 depletion. hTERT-RPE1 cells were treated with siRNAs (C-E) or in vitro-diced esiKi-67 (F-H) for 72 hrs. Panels C, F: Cells were analyzed by immuno-RNA-FISH to localize XIST (green) and macroH2A (red). Scale bar: 10 μm. Panels D, G: Quantitation of cells that displayed reduced macroH2A staining is shown. Total cells assayed= 248 for si-scramble, 291 for siKi-67, 218 for esi-luciferase, 236 for esiKi-67. Panels E, H: immunoblot analyses of Ki-67 depletions.

### Ki-67 affects the S phase nucleolar association of the inactive X chromosome

The perinucleolar space is a subset of the heterochromatic compartment; another frequent location for heterochromatin is at the nuclear periphery, adjacent to the nuclear lamina (49). Accordingly, the Xi is usually localized to one of these two preferred locations (24). However, heterochromatic sequences can dynamically relocalize from nucleoli to the periphery, either during cell division or upon perturbing the nucleolus with actinomycin (49-52). Because Ki-67 depletion affected heterochromatic marks only on Xi chromosomes away from the nuclear periphery (Figs. 10-13), we hypothesized that it might also affect the interphase localization of the Xi. We first examined Xi localization in asynchronous hTERT-RPE1 cells, using immuno-RNA-FISH to detect Xi-associated lncRNA XIST and nucleolar protein fibrillarin. Indeed, Ki-67 depletion resulted in a partial but statistically significant reduction in Xi-nucleolar associations (Figure 15A-D). This loss of nucleolar association was accompanied by an increase of similar magnitude in Xi-lamina associations, and similar results were observed with our two distinct Ki-67 depletion reagents (Figures 15B, D). However, in 293T cells, we observed no significant alteration in Xi-nucleolar associations (Fig. 15E-H). Thus, distribution of the Xi within the interphase nucleus is sensitive to Ki-67 depletion in a cell type that induces p21.

**Figure 15.**
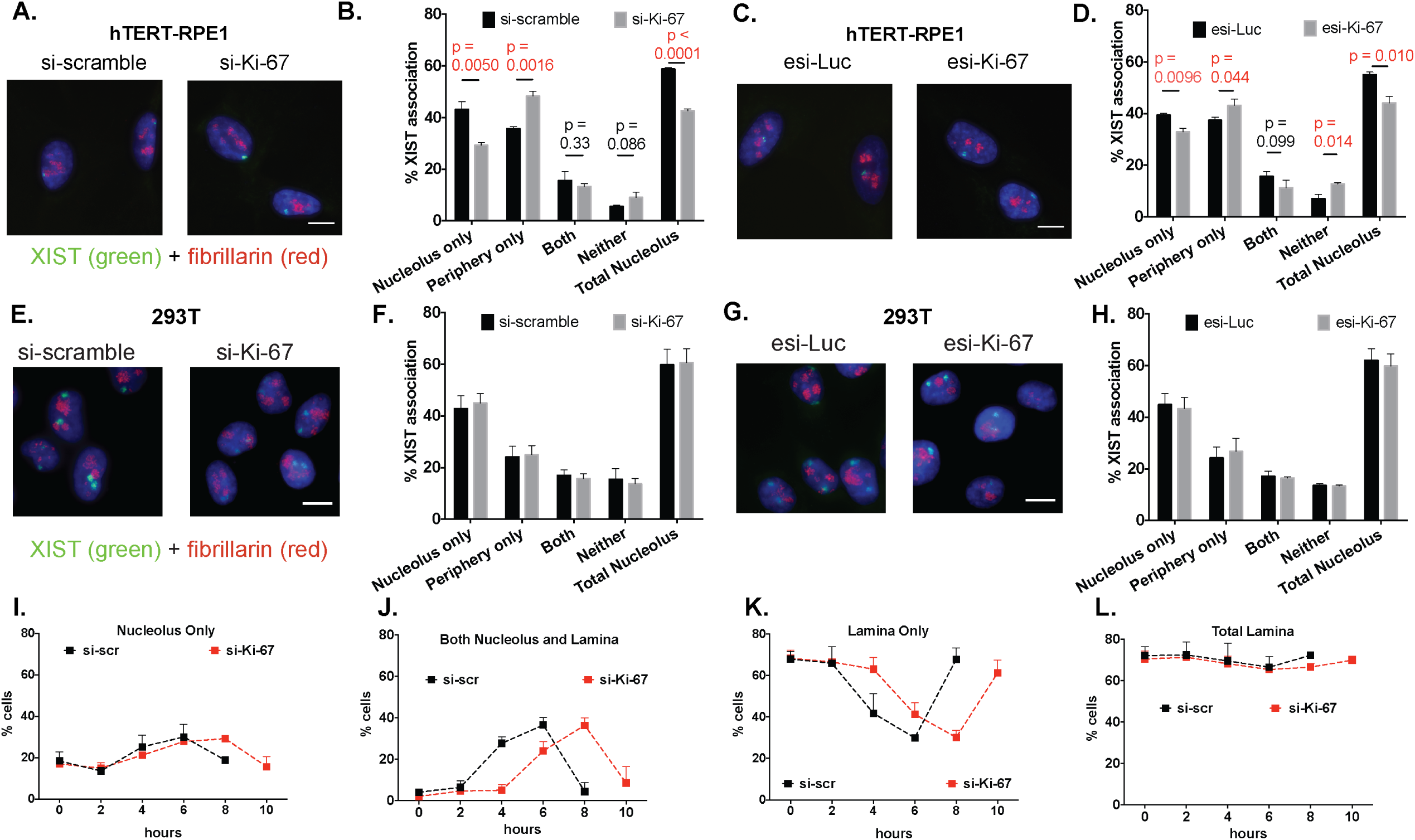
Depletion of Ki-67 redistributed the Xi chromosome within hTERT-RPE1 but not 293T nuclei. Scale bars, 10 μm. A. Fluorescence microscopy images of representative hTERT-RPE1 cells treated either with scramble control or Ki-67-targeted siRNAs as indicated. Cells were analyzed by RNA-FISH to detect XIST RNA (green) marking the Xi, and by immunofluorescence with anti-fibrillarin antibodies (red) to label nucleoli. DNA was stained with DAPI (blue). B. Quantification of XIST association frequencies from panel A experiments. XIST associations with the indicated locations were counted; “total nucleolar” indicates the sum of XIST signals that are exclusively nucleolar plus those that also are on the nuclear periphery. Three biological replicate experiments were performed with mean percentage association and SDs graphed. Total cells assayed = 363 for si-scramble, 376 for siKi-67. Holm-Sidak corrected p-values comparing si-scramble and siKi-67 treatments are indicated, with p-values < 0.05 in red. C. Fluorescence microscopy images of representative hTERT-RPE1 cells treated either with luciferase- or Ki-67-targeted esiRNAs as indicated. Cells were stained as in panel A. D. Quantification of association frequencies from panel C experiments. Total cells assayed = 348 for esi-luciferase, 391 for esiKi-67. E. Fluorescence microscopy images of representative 293T cells treated either with scramble control or Ki-67-targeted siRNAs. F. Quantification of XIST association frequencies from panel E experiments. Total alleles assayed = 272 for si-scramble, 298 for siKi-67. Holm-Sidak adjusted p-values comparing association frequencies were > 0.97 for all comparisons. G. Fluorescence microscopy images of representative 293T cells as panel E, except that cells were treated with in vitro-diced esiRNAs as depletion reagents. H. Quantitation of XIST association frequencies from panel G experiments. Total alleles assayed = 250 for esi-luciferase, 270 for esiKi-67. Holm-Sidak adjusted p-values comparing association frequencies were > 0.98 for all comparisons. Panels I-L: Frequencies of Xi associations versus time. All associations were measured in HU-synchronized cells as in Fig. 7A-B, with the averages and std. dev. from 3 independent experiments. At each time point, > 300 cells were counted. I. Xi-nucleolar only associations (Xi chromosomes associated with nucleoli but not lamina). J. Xi chromosomes associated with both the lamina and nucleoli simultaneously. K. Xi-lamina only associations (Xi chromosomes associated with lamina but not nucleoli). Note that Xi-nucleolar associations (panels I-J) peak and Xi-lamina only (panel K) associations reach a minimum when the majority of cells are in mid-late S phase, which is delayed in Ki67-depleted cells by 2 hr. L. Xi-total laminar associations (Xi chromosomes associated either with the lamina or with both lamina and nucleoli simultaneously).

Previous studies in mouse cells showed that the Xi-nucleolar association is cell-cycle regulated, occurring most prevalently in S phase cells (24). Therefore, we examined Xi localization in hTERT-RPE1 cells prepared in the same manner as in the cell synchronization experiments in Figure 7A-B. Consistent with published data from mouse cells (24), the frequency of Xi-nucleolar associations peaked in middle S-late S phase transition; this was true for the frequency of Xi associations that were exclusively at the nucleolus (Fig. 15I), and for the frequency of Xi chromosomes simultaneously associated with both the nucleolus and the lamina (Fig. 15J). These peaks occurred in both the control and Ki-67-depleted populations, with the Xi-nucleolar interaction peaks delayed two hours in the latter case. The two-hour shift correlates with the delay in S phase entry in the Ki-67-depleted cells (Fig. 7).

As suggested by the Xi localization data from asynchronous cells (Fig. 15A-D), Xi associations with the nucleoli and lamina were inversely related, so that the peak of nucleolar associations (Figure 15I-J) coincided with the lowest frequencies of laminar associations (Fig. 15K). We note that when the total Xi-laminar association frequencies were counted by summing the exclusively laminar associations (Fig. 15K) plus those also associated with nucleoli simultaneously (Fig. 15J), we observed little change during the experiment (Figure 15L). Thus, the biggest changes during S phase are lamina-associated Xi becoming transiently also associated with nucleoli (e.g. compare Fig. 15J and 15K). Together, these data indicated that cell cycle-regulated Xi-nucleolar associations are delayed in concert with DNA synthesis upon depletion of Ki-67 in hTERT-RPE1 cells. Thus, checkpoint activation upon Ki-67 depletion affects cell cycle progression and gene expression, and these effects are correlated with altered Xi heterochromatin in female hTERT-RPE1 cells.

## Discussion

### Cell-type specific responses to Ki-67 depletion

Our studies show that Ki-67 expression is important for normal S phase progression in primary human foreskin fibroblasts, non-transformed fibroblast lines (WI-38 and IMR90), hTERT-immortalized BJ fibroblasts, and hTERT-immortalized RPE1 cells (23), a human female retinal pigment epithelial cell line with a diploid karyotype (53). In contrast, in cancer-derived 293T, U2OS and HeLa cells, depletion of Ki-67 did not cause defects in cell proliferation. Therefore, we conclude that Ki-67 is required for normal cell cycle progression in some but not all cell lines. Our data distinguish two types of responses to depletion of Ki-67 in human cells, depending on whether cells are able to mount a p21-dependent checkpoint.

hTERT-BJ fibroblasts were sensitive to Ki-67 depletion in our assays; however, a previous study indicated that shRNA-mediated Ki-67 depletion did not affect hTERT-BJ cell cycle re-entry after starvation (12). Therefore, not all assays can detect the effects of Ki-67 depletion. For example, two assays used in the previous study are insensitive to the cell cycle progression delays that we observe upon Ki-67 depletion. First, one-dimensional flow cytometry of propidium iodide-stained asynchronous populations cannot detect the S phase delays we detect in BrdU labeling experiments; second, a 3 hour EdU pulse is too long to capture the 2-hour S phase entry delay. Together, our data indicate that the effects of Ki-67 on S phase progression are transient in sensitive cell types and therefore most easily observed using short pulses of labeled deoxynucleotides.

### Characteristics of checkpoint activation caused by Ki-67 depletion

Meta-analyses of our RNAseq data showed that Ki-67 depletion in hTERT-RPE1 cells resulted in frequent repression of Rb/E2F-regulated, G1/S expressed genes. However, depletion of the Rb checkpoint protein only partially relieved transcriptional repression and did not restore normal percentages of S phase cells, indicating that additional factors were responsible for the altered cell cycle profile in Ki-67 depleted cells.

A strong candidate for such a factor is the DREAM complex, which represses G1/S-expressed cell cycle genes in a p21-dependent manner (26, 54). We noticed frequent downregulation of DREAM complex targets in Ki-67-depleted cells, which lead us to test whether sensitivity to Ki-67 depletion was also p21-dependent. Notably, both the transcriptional alterations and cell cycle perturbations caused by Ki-67 depletion were partially relieved by simultaneous depletion of p21. Importantly, cell lines that induce p21 upon Ki-67 depletion are those that inhibit transcription of DNA replication genes and delay entry into S-phase. Thus, this study implicates a p21-dependent checkpoint in cells sensitive to Ki-67 depletion.

A recent study shows that PD0332991, a small molecule CDK4/CDK6 inhibitor (CDKi), depletes Ki-67 protein levels in some but not all cell lines (55). In “CDKi-sensitive” cells, this compound causes G1 cell cycle arrest via an Rb-mediated checkpoint, inhibiting Ki-67 and cyclin A gene transcription while proteasome-mediated degradation destroys existing Ki-67 protein molecules. CDKi-sensitivity appears similar to many of the responses we observe upon Ki-67 depletion, and CDKi-sensitive cells include IMR90 and primary fibroblasts that we found to be “Ki-67 sensitive”. MCF7 breast adenocarcinoma cells are also CDKi-sensitive (55), and we find that upon Ki-67 depletion MCF7 cells induce p21 and down-regulate DNA replication genes. In contrast, HeLa and U2OS cells are not sensitive to either CDKi or Ki-67 depletion. Thus, depletion of Ki-67 via CDKi treatment (55) or via siRNA (this work) often leads to similar outcomes.

However, there is a counterexample to these correlations. CDKi treatment blocks S phase entry and depletes Ki-67 in MDA-MB-231 breast adenocarcinoma cells (55). In contrast, upon Ki-67 depletion, MDA-MB-231 cells did not display either p21 induction or transcriptional down-regulation of S phase genes. Therefore, proteasome-mediated degradation of Ki-67 via CDK4/6 inhibition is not equivalent to siRNA-mediated Ki67 depletion in all cell types. We hypothesize that a key difference is related to induction of p21 in “Ki-67-sensitive” cell lines. p21 contributes to G1/S arrest via multiple mechanisms. As a CDK inhibitor (32,33), p21 blocks CDK-mediated Rb phosphorylation thereby inhibiting E2F-driven transcription(56). Likewise, it maintains activity of the transcriptionally repressive DREAM complex which contain Rb-related p107/p130 “pocket protein” subunits (26,27,54). p21 also directly interacts with PCNA and directly inhibits DNA synthesis (35). Therefore, the lack of p21 induction in MDA-MD-231 cells may be the key factor explaining the lack of “Ki-67 sensitivity” in this cell line. Future experiments will be required to determine whether the activation of the DREAM complex or direct inhibition of DNA synthesis machinery is more important for the “Ki-67-sensitive” phenotype associated with p21 induction.

Regarding the defect in p21 induction in MDA-MB-231 cells, we note that they express a gain-of-function R280K allele, which dominantly blocks p21 induction (38). Thus, p53 status is likely a critical aspect of the different cell cycle responses to Ki-67 depletion in many cell lines. However, sensitivity to Ki-67 depletion cannot always be predicted strictly by p53 status. For example, HCT116 cells express wild-type p53 (38) but are not sensitive to the CDKi PD0332991 (55), or to Ki-67 depletion (Fig. 9J). Because Ki-67 expression predicts the differential response of different cell lines to CDKi treatment during xenograft tumor formation (55), understanding how different checkpoint mutations alter Ki-67 expression and sensitivity to its depletion are important goals for developing stratified approaches to cancer therapies.

### Ki-67 contributes to the interphase localization of the Xi chromosome

Nucleoli are non-membrane bound organelles within the nucleus. Not only are these sites of synthesis and assembly of ribosome components, the periphery of these organelles plays an important role in higher order chromosome localization (49,57). Specifically, the nucleolar periphery houses a subset of the cellular heterochromatin, which exchanges dynamically with lamina-associated heterochromatin (50-52). Like other heterochromatin regions, high resolution analysis of nucleolus-associated domains (termed NADs) in human cells reveals enrichment of satellite repetitive DNAs and repressive histone marks (51,58,59). Major questions in chromosome biology are how heterochromatin regions are partitioned to different intranuclear locations, and how these interactions are governed by cell cycle progression.

As a region of facultative heterochromatin, the Xi chromosome in female cells is enriched in NAD loci, usually localized to either the nucleolar periphery or to the nuclear lamina (24,40,58). In mouse cells, the Xi-nucleolar association is cell cycle-dependent, with frequencies peaking during mid-to-late S phase. A genetic deletion was used to show that the long non-coding RNA Xist, which is expressed from the Xi chromosome, is required for normal Xi-perinucleolar localization. Deletion of Xist results in diminished H3K27me3 enrichment and increased synthesis of Cot-1-hybridizing, repeat-derived RNAs on the Xi (24). These data suggest that perinucleolar localization of Xi contributes to the maintenance of heterochromatin structure. More recently, depletion of long non-coding RNA Firre was shown to reduce association of the Xi to nucleolus in mouse cells, and also reduces H3K27me3 density on the Xi (60). However, *Firre* depletion has minimal effects on Xi gene silencing, consistent with the idea that multiple functionally overlapping factors affect Xi heterochromatin localization and gene silencing.

Here, we discovered that Ki-67 affects the Xi-nucleolus interaction. Analysis of synchronized RPE-1 cells shows that the association appears more slowly in Ki-67-depleted cells, coincident with the delay in S phase entry. In addition to this delay, Ki-67 depletion alters some of the heterochromatin characteristics of the Xi, causing significantly increased levels of Cot-1-hybridizing RNAs and Pol ll, and decreased enrichment of H3K27me3 and H4K20me1. This loss of heterochromatic properties is partial in the cell population, and we do not detect uniform reactivation of Pol II genes on the Xi. These data are consistent with previous studies showing that multiple overlapping mechanisms maintain the inactive status of the Xi (60,61). We note that XIST levels are not reduced upon Ki-67 depletion (Supplemental Table 2; Fig. 10), suggesting that altered XIST levels are unlikely to explain the effect of Ki-67 on the Xi. Instead, our data is consistent with the view that Ki-67 is one of the factors that contributes to the maintenance of heterochromatic structures of the Xi (12) in this case in a manner coupled to S phase progression.

Recent data show that the nuclear lamina localization mediated by the interaction between XIST and the lamina B receptor facilitates the spreading of XIST on the Xi chromosome, which in turn contributes to transcriptional silencing (62). This raises the question of whether there are specific protein factors that contribute to the association of the Xi with the nucleolus in addition to the lncRNAs XIST and FIRRE (24,60). Could Ki-67 be such a factor? In support of this idea, Ki-67 is also required for association of other heterochromatic regions with nucleoli in interphase cells (11,12,22).

It appears that the erosion of heterochromatic features on the Xi occurs in a significant fraction of Ki-67-depleted cells away from the nuclear lamina, but the lamin-associated Xi chromosomes are not altered. There are two possibilities to explain these data. First, it may be that lamina association confers protection from heterochromatin changes. Alternatively, Xi chromosomes that are most severely affected by Ki-67 depletion may preferentially relocalize away from the lamina. Because our shRNA-based experiments necessitate a 72-hour period to achieve strong Ki-67 depletion, there is likely passage through multiple mitoses for each cell during this period. As Ki-67 is a key component of the perichromosomal layer that envelopes each mitotic chromosome (11,15), it is tempting to speculate that the loss of Ki-67 affects the reassociation of heterochromatic sequences with the nuclear lamina or nucleoli after mitotic exit.

## Materials and Methods

### Antibodies and Immunoblotting

The following antibodies were used in this work:

rabbit anti-Ki-67 (Abcam Ab15580)

mouse anti-beta-tubulin (Ubpbio Y1060)

rabbit anti-fibrillarin (Abcam ab5821-100)

mouse anti-BrdU antibody (MoBu-1) (Abcam ab8039)

mouse anti-p21 antibody (Abcam ab109520)

rabbit anti-mcroH2A.1 (Abcam ab37264)

mouse anti-Rb antibody (4H1) (Cell signaling 9309)

mouse anti-H4K20Me1 (Active motif 39727)

mouse anti-RNA polymerase II, clone CTD4H8 (Millipore 05-623)

mouse anti-nucleophosmin (Santa Cruz Biotechnology sc-32256)

rabbit anti-WSTF (Cell signaling 2152)

rabbit anti-H3K27Me3 (Active motif 39535)

Amersham ECL Rabbit IgG, HRP-linked whole Ab (GElifescience NA934)

Donkey anti-Rabbit IgG (H+L) Secondary Antibody, Alexa Fluor® 594 (Life Technologies A-21207)

Donkey anti-Rabbit IgG (H+L) Secondary Antibody, Alexa Fluor® 488 (Life Technologies A-21206)

Streptavidin, Alexa Fluor® 488 Conjugate (Invitrogen S-32354)

DyLight 594 Labeled Anti-Digoxigenin/Digoxin (Vector Labs DI-7594)

For immunoblotting, cells were collected 3 days after RNAi transfection. Whole-cell lysates were extracted in 20 mM Tris-HCl 7.5, 1% SDS and 10% glycerol supplemented with protease inhibitor cocktail (Sigma P8340-1ml). The lysates were sonicated in a Bioruptor set on high power for one 5 min cycle, with 30s on/30s off. 15 ⍰g of each lysate were separated by SDS-PAGE, transferred to PVDF membrane and probed as described in the figure legends.

### Cell Cultures

hTERT-RPE1 cells (a kind gift from Dr. Judith Sharp) and hTERT-BJ were cultured in DMEM-F12 medium (VWR 12001-600) with 10% fetal bovine serum (FBS, Hyclone #SH30910.03), 1% penicillin/streptomycin, 5% L-glutamine and 7.5% sodium bicarbonate solution.

HeLa and U2OS cells were propagated in DMEM medium supplemented with 10% fetal bovine serum. Cells were maintained > 25% confluence and passaged every three days. Human foreskin fibroblasts (HFF, a kind gift from Dr. Jennifer Benanti, University of Massachusetts Medical School (63) were maintained in DMEM containing 10% FBS and antibiotic/antimycotic solution. HFF cells were grown at > 25% confluence and were split 1:4 every 2 days. All cells were maintained in a 37 ^°^C incubator with 5% CO_2_WI-38 cells were maintained in DMEM and supplemented with 10% FBS, 2 mM L-glutamine and antibiotic/antimycotic solution (Life Technologies, Carlsbad, CA). WI-38 cells were grown at > 25% confluence and were split 1:4 every 2 days.

IMR-90 and MDA-MB-231 were maintained in DMEM medium with 10% FBS, 2mM L-glutamine, and antibiotic/antimycotic solution. The cells were cultured at > 25% confluence and were split every 2 days.

HCT 116 (a kind gift of Dr. Anastassiia Vertii, University of Massachusetts Medical School) and 293T cells were cultured in DMEM medium with 10% FBS. The cells were split 1:4 every 3 days

MCF7 cells were grown in RPMI medium with 10% FBS. The cells were split 1:3 every 3 days.

For hydroxyurea treatment, hTERT-RPE1 cells were cultured in the presence of 2 mM hydroxyurea for 15 h, then washed with three times phosphate-buffered saline (PBS) and released into hydroxyurea-free medium and harvested at the indicated time points.

### sgRNA design

For editing the siRNA target site in the endogenous Ki-67 locus, four highest scoring single guide RNAs (sgRNAs) targeting nucleotides 9661-9722 of the genomic DNA (NG_047061) were selected by using the CRISPR Design web tool at http://crispr.mit.edu/ (64). The sgRNA sequences are shown below:

5’-ACGTGCTGGCTCCTGTAAGT-3’ (antisense)

5’-TCTAGCTTCTCTTCTGACCC-3’ (sense)

5’-GATCTTGAGACACGACGTGC-3’ (antisense)

5’-CTTCTGACCCTGGTGAGTAG-3’ (sense)

These were cloned into a variant of the pX330 plasmid (64) with a puromycin-resistance cassette (a kind gift from Kurtis McCannell and Dr. Thomas Fazzio, University of Massachusetts Medical School) as previously described (65). To determine the most efficient sgRNAs, 293T cells were transfected with FuGENE HD (Promega, catalog number E2311) according to manufacturer’s instructions. 500 ng of sequence-verified CRISPR plasmid (pSpCas9-sgRNA) was transfected into 200 × 10^3^ cells in 24 well dish. 48 hours post-transfection, DNA was extracted using QuickExtract DNA extraction solution (Epicentre, catalog number QE09050) according to manufacturer’s instructions. Genomic DNA was PCR amplified using a *Taq* DNA Polymerase (New England Biolabs, catalog number M0273). The following primers were used for PCR amplification:

F2 primer: GGGTTCCAGCAATTCTCCTG

R primer: TCACCAAGGGAAAGTTAGGC

514 bp PCR products were ran on agarose gel and purified using Zymoclean Gel DNA recovery kit (Zymo Research, catalog number D4007) and sent for Sanger sequencing at Genewiz sequencing facility. The following primer was used for sequencing:

F primer: GCCAGGCTGTTCTCAAACTC

To assess gene editing in 293T cells by four sgRNA plasmids, TIDE web tool at <u>https://tide-calculator.nki.nl/</u> was used (66). Trace data for PCR fragment from GFP-transfected 293T cells were used as a control sample chromatogram, and trace data for PCR fragments from CRISPR plasmid-transfected cells were used as test sample chromatograms. The following sgRNA was selected due to its efficiency in cutting and proximity to the siRNA site:

5’-GATCTTGAGACACGACGTGC-3’ (referred to as sgKi-67 from now on).

### Co-transfection of CRISPR plasmid and HDR template into hTERT-RPE1 cells

An HDR repair template carrying the siRNA-resistance conferring mutations, EcoO109I site and 700 bp homology flanks on each side was purchased as a gBlock from Integrated DNA Technologies. The template was cloned into pCR2.1 (Thermo Fisher Scientific, catalog number K200001) according to manufacturer’s instructions.

The sgKi-67 was cloned into a variant of the pX330 plasmid(64) with a neomycin-resistance cassette (a kind gift from Kurtis McCannell and Dr. Thomas Fazzio, University of Massachusetts Medical School) to facilitate gene editing in hTERT-RPE1. 3.3 μg of a 1:1 (v/v) mix of repair template and CRISPR plasmid were transfected using FuGENE HD into 75 ×10^3^ cells in 6 well dish. Starting 48 hours post-transfection, cells were cultured in selection medium with 800 μg/ml of G418 (Sigma-Aldrich, catalog number A1720) for 7 days, with selection medium being changed every other day. Cells were recovered in G418-free medium for 4 days, after which cells were trypsinized and diluted to 0.5 cells per 200 μl and seeded into 96-well plates. A week later plates were inspected for wells with single colonies, and 4 days after that replica plated into 2× 96-well plates. One plate was frozen down, the second one used to maintain and passage the cells. Once cells on the third plate were at least ∼70% confluent, DNA was extracted using QuickExtract DNA extraction solution and PCR amplified using the following primers.

F3 primer: TGGCCCATTTATGAGAAAACTGA

R2 primer: GGGAACAGACTTCAATTCTCCA

1523 bp PCR products were further digested with EcoO109I restriction enzyme (New England Biolabs, catalog number R0503S). PCR products from successfully integrated clones were expected to be digested at 751 and 772 bp. PCR products for clones positive for the EcoO109I digested bands were ran on agarose gel, purified using Zymoclean Gel DNA recovery kit and sent to Genewiz for Sanger sequencing. Primers F2 and R2 shown above were used for Sanger sequencing.

### Immunofluorescence

Cells grown on glass coverslips were fixed in 4% paraformaldehyde for 10 min and then permeabilized with 0.5% Triton X-100 for 10 min at room temperature. The fixed cells were blocked in 5 % goat serum for 30 min, and incubated in primary antibody at 37 ^°^C in a humidified chamber for 1h. The cells were washed with PBS for 5 min three times, incubated with secondary antibody for 1 h at 37 ^°^C in humidified chamber, followed by three PBS washes, 5 min each. Slides were then incubated with 130 ng/ml 4,6-diamidino-2-phenylindole (DAPI) for 5 min and mounted in Vectashield mounting medium (Vector Lab, H-1000)

Images were taken on a Zeiss Axioplan2 microscope with a 63× objective. Entire cells were imaged via Z stacks taken at 200 nm step-intervals. Approximately 25 stacks were taken per cell, and displayed as 2D maximum intensity projections generated using AxioVision version 4.6.

The Xi-nucleolar association frequencies were scored in a blinded manner. Entire cells were imaged via Z stacks taken at 200 nm step-intervals. Approximately 25 stacks were taken per cell, and displayed as 2D maximum intensity projections generated using AxioVision version 4.6. The Xi-nucleolar association frequencies on individual coverslips were scored in a blinded manner. The criteria for Xi-nucleolar or lamina association was that there were no pixels between the fluorescence signals from the XIST FISH probe and fibrillarin immunostaining (for nucleolar association) or DAPI staining of the nuclear edge (for lamina association). Densitometry of individual immunostained cells was performed in Image J 10.2 (48), using the macro script of the RGB Profiles Tool for all experiments. The quantifications of H3K27me3, macroH2.A and H4K20me1, Cot-1 and Pol II enrichment were also performed in Image J 10.2 (48).

### Visualization of 5-ethynyl-2-deoxyuridine(EdU)-labeled nascent DNA

hTERT-RPE1 cells were grown on glass coverslips in DMEM/F12 media as described above. 5-ethynyl-2-deoxyuridine (EdU) was added to the culture medium at concentration of 10 μM for 20 min. After labeling, cells were washed three times with PBS. Cells were then permeablized in 0.5% Triton X-100 for 30 seconds and then fixed in 10% formaldehyde for 10 min. Cells were then rinsed twice with PBS and then incubated 30 min in 100 mM Tris-HCl pH 8.5, 1 mM CuSO_4_, 100 mM ascorbic acid plus 50 mM carboxyrhodamine 110-azide for click-chemistry labeling. After staining, the cells on coverslips were washed three times with PBS plus 0.5% Triton X-100, 5 min each. Cells were then counterstained with DAPI, mounted in Vectashield and imaged by fluorescence microscopy as above.

### Immuno-RNA-FISH and RNA-FISH

The plasmid pGIA which contains human XIST exons 4, 5 and 6 was a gift from Dr Judith Sharp. Cot-1 probe was from Invitrogen (Sigma 11581074001). The probes were labeled either with biotin-14-dCTP or digoxigenin-11-dUTP using the BioPrime DNA labeling system (Invitrogen 18094-011).

For immunofluorescence (IF) combined with RNA-FISH, IF was performed first as above. Cells were then re-fixed in 4% paraformaldehyde for 10 min at room temperature. The cells were then dehydrated in 75%, 85%, 95% and 100% ethanol for 2 min each.

Approximately 150 ng of each probe was mixed with 20 μg single-stranded salmon sperm DNA (Sigma-Aldrich) and 12 μg E. coli tRNA and then air-dried in a speed vacuum, resuspended in 20 ⍰l 50% formamide / 50% hybridization buffer (20% Dextran Sulfate in 4× SSC), denatured at 80 ^°^C for 10 min and pre-annealed at 37 ^°^C for 30 min prior to hybridization. Hybridizations were performed overnight in a humidified chamber at 37^°^C. The next day, cells were washed for 20 min in 50% formamide in 2× SSC at 37°C and then for 20 min in 2× SSC at 37°C and 20 min in 1× SSC at 37°C. The hybridized probes were detected by incubation with either Alexa fluor-488 conjugated to streptavidin (Invitrogen S-32354) or Dylight 594 labeled anti-Digoxigenin/Digoxin (Vector Labs, DI-7594) at 1:500 dilutions for 60 min in a 37^°^C humid chamber. After incubation, slides were washed twice in 50% formamide, 2× SSC for 5 min and once in 1× SSC for 5 min in a 37^°^C humid chamber before DAPI staining as above.

### RNAi experiments

The siRNA targeting human Ki-67 is from the collection of Silencer Select Predesigned siRNAs (Thermo Fisher Scientific), and targets nucleotides 559-577 of the cDNA (NM_002417.4).

Its sequence is as follows:

sense CGUCGUGUCUCAAGAUCUAtt,

antisense UAGAUCUUGAGACACGACGtg

TP53 (NM_00546.5)

Forward primer for hTP53: GAAAUUUGCGUGUGGAGUAtt

Reverse primer for hTP53:UACUCCACACGCAAAUUUCct

p21(NM_078467.2)

Forward primer for hp21: CAAGGAGUCAGACAUUUUAtt

Reverse primer for hp21: UAAAAUGUCUGACUCCUUGtt

esiRNA targeting human Ki-67 was generated by in vitro RNaseIII cleavage of T7 RNA polymerase-synthesized transcripts, as previously described (21,67), and targets internal repeat regions at nucleotides 3611-4047, 3979-4357, 4705-5098 and 6913-7347 of the cDNA (NM_002417.4).

Forward primer for hKi-67

gcgtaatacgactcactataggGTGCTGCCGGTTAAGTTCTCT

Reverse primer for hKi-67

gcgtaatacgactcactataggGCTCCAACAAGCACAAAGCAA

Forward primer for luciferase

gcgtaatacgactcactataggAACAATTGCTTTTACAGATGC

Reverse primer for luciferase

gcgtaatacgactcactataggAGGCAGACCAGTAGATCC

Cells were transfected with Lipofectamine RNAi MAX (Invitrogen Catalog number 13778100) following manufacturer’s instructions.

For esiRNA transfection, 500ng of esiRNA targeting either luciferase control or Ki-67 was transfected into 40 × 10^3^ cells in a 6-well dish.

For siRNA transfection, 40 nM of either scramble or siKi-67 was transfected into 40 × 10^3^ cells in a 6-well dish.

The cells were harvested 72 hrs after transfection for immunoblotting, RT-qPCR, RNA-seq, FACS or FISH analysis.

### Flow cytometry

BrdU incorporation was analyzed based on published prototcols (68). Cells were pulsed labeled with 50 μM bromodexoyuridine (BrdU) for the indicated time periods. Cells were then washed twice with PBS and fixed in 70% ethanol at 4^°^C for 1 hour. Post-fixed cells were denatured in 2N HCl/0.5% Triton-X for 30 minutes. After denaturation, cells were washed once in 0.1 M sodium tetraborate for 2 minutes and once in PBS/1% BSA. After that, cells were resuspended in 1 μg/ml anti-BrdU antibody/PBS/1% BSA for 1 hr, followed by three washes with 0.5% Tween 20/1% BSA/PBS. The cells were incubated with 0.5 μg/ml secondary antibody/PBS/1% BSA for 30 minutes and counterstained with 50 μg/ml propidium iodide / PBS and analyzed on a LSR II (BD Biosciences). The data was analyzed with FlowJo v9.9.4 software (TreeStar, Ashland, OR).

### RNA isolation and real time quantitative PCR

Total RNA from cells 72 hours post-transfection was isolated using Trizol (Invitrogen 15596026) following manufacturer’s instructions and purified using the RNeasy kit (Qiagen 74104).

One microgram of RNA was subjected to reverse transcription with SuperScript II Reverse Transcriptase (Invitrogen 18064014). qPCR reactions were performed on an Applied Biosystem StepOnePlus machine (Life Technologies) using FAST SYBR mix (KAPA Biosystem). The program used is as follows: hold 98°C for 30 s, followed by 40 cycles of 95°C for 10 s and 60°C for 30 s. All the signals were normalized with beta-actin as indicated in the figure legends and the 2^-ΔΔCt^ method was used for quantification (Life technologies). Primer sequences are designed by Primer3Plus software (69). All oligonucleotides for qPCR are listed in Supplemental Table 5.

### RNAseq: sample preparation and analysis

RNA was isolated as described above. Libraries from two replicates for each condition were constructed in a strand-specific manner via the dUTP method by BGI and sequenced using Illumina-HiSeq 2000/2500 platform (BGI) as single-end 50-base reads. 29M and 31M mapped reads were obtained from two si-scramble-treated controls, 28M and 29M were obtained from two siKi-67 knockdown replicates.

Reads were aligned to the human reference genome (hg19) using Tophat 2.0.14 (70,71). Differential expression analysis was determined by Cufflinks 2.2.1 (72). In addition, the Reactome analyses were performed using Bioconductor package ChIPpeakAnno (version 3.2.0)(73,74). Genes that showed differential expression with BH-adjusted q value < 0.05 (75) between control and Ki-67 depletion samples were selected for the Reactome analysis.

For the SNP analysis, the genotype (SNPs) information of the RPE cell line (76) from GEO sample GSM1848919. (https://www.ncbi.nlm.nih.gov/geo/query/acc.cgi?acc=GSM1848919) was annotated based on R bioconductor package "SNPlocs.Hsapiens.dbSNP144.GRCh38".

*SNPlocs.Hsapiens.dbSNP144.GRCh38: SNP locations for Homo sapiens (dbSNP Build 144)*. R package version 0.99.20.) The SNP locations were further annotated by ChipPeakAnno package (73).

### Accession Numbers

The data discussed in this publication have been deposited in NCBI’s Sequence Read Archive (SRA, http://www.ncbi.nlm.nih.gov/sra/) and are accessible through SRA Series accession number SRR4252548.

## Acknowledgments

We would like to thank Michael Brodsky for the generous use of the AxioPlan microscope. This work was supported by NIH grants R01 GM55712 and U01 DA040588 to PDK.

## Legends for Supplemental tables

**Supplemental Table 1.** Top reactomes detected in siKi-67 RNAseq data.

Reactome analysis of transcriptional changes reveals functional grouping of pathways altered upon Ki67 depletion in hTERT-RPE1 cells. Table shows all pathways enriched with p value < 5 × 10^-5^. Log2FC value for selected DNA replication-related genes are shown at the bottom

**Supplemental Table 2.** Complete RNA-seq data from siRNA-treated hTERT-RPE1 cells.

Each transcript that mapped uniquely to the genome is listed. Value 1 is the mean FPKM from two biological replicates in si-scramble-treated hTERT-RPE1 cells. Value 2 is the mean FPKM from two biological replicates in siKi-67-treated hTERT-RPE1 cells. Fold change is the ratio of value 2 over value1 in log2 range. Status, Test_stat, p_value and q_ value are generated from Cufflinks 2.2.1(1). Status: “OK” indicates a successful test, “NOTEST” indicates insufficient alignments for testing. Test_stat: displays the significance of the observed change in FPKM. p_value: uncorrected p-values of the test statistic. q_value: p-value adjusted by the false discovery rate (FDR). The criteria of significance is defined as q < 0.05.

**Supplemental Table 3.**

Comparison of our si-Ki-67 RNAseq data to meta-analyses of cell cycle regulators (2). Separate sheets for the Cell cycle, E2F, and DREAM data are included. Cell cycle: The “Adjusted Cell Cycle Score” equals the Cell cycle expression score (Negative for G1/S-expressed genes, positive for G2/M-expressed genes, Table S10 in (2)), + 1 point for regulation by G2/M regulator MMB/FOXM1, -1 point for regulation by G1/S regulators Rb/E2F. “si-Ki-67 log2FC” and “q-values” are from the RNAseq analysis of hTERT-RPE1 cells (Figure 1). Genes with the same Adjusted Cell Cycle score were analyzed in the same bin. “Significant upregulation” was TRUE if the log2FC> 0 AND q< 0.05. “Significant downregulation” was TRUE if log2FC< 0 AND q< 0.05. Benjamini-adjusted p-values (3) for these enrichments were calculated and graphed in Figure 8.

E2F: The Rb/E2F binding scores come from Table S9 of (2).

DREAM: The DREAM binding scores come from Table S7 of (2).

**Supplemental Table 4.** Analysis of X-linked SNPs in the RNAseq data.

The RNAseq analyses of hTERT-RPE1 cells were compared to SNPs from the genome sequence these cells (4). We also compared these data to previous analysis of escapers from Xi silencing (5, 6).

**Supplemental Table 5.** Oligonucleotides used in this study

Table shows primers used for RT-PCR or for generating esiRNA.

## References

1. Gerdes J, Schwab U, Lemke H, Stein H. 1983. Production of a mouse monoclonal antibody reactive with a human nuclear antigen associated with cell proliferation. Int J Cancer 31:13–20.2.

2. Gerdes J, Lemke H, Baisch H, Wacker HH, Schwab U, Stein H. 1984. Cell cycle analysis of a cell proliferation-associated human nuclear antigen defined by the monoclonal antibody Ki-67. J Immunol 133:1710–1715.

3. Gerdes J, Stein H, Pileri S, Mt R, Gobbi M, Ralfkiaer E, Km N, Pallesen G, Bartels H, Palestro G. 1987. Prognostic relevance of tumour-cell growth fraction in malignant non-Hodgkin's lymphomas. Lancet 2:448–449.

4. Dowsett M, Nielsen TO, A'Hern R, Bartlett J, Coombes RC, Cuzick J, Ellis M, Henry NL, Hugh JC, Lively T, McShane L, Paik S, Penault-Llorca F, Prudkin L, Regan M, Salter J, Sotiriou C, Smith IE, Viale G, Zujewski JA, Hayes DF. 2011. Assessment of Ki67 in Breast Cancer: Recommendations from the international Ki67 in breast cancer working Group. J Natl Cancer Inst 103:1656–1664.

5. Verheijen R, Kuijpers HJ, Schlingemann RO, Boehmer AL, van Driel R, Brakenhoff GJ, Ramaekers FC. 1989. Ki-67 detects a nuclear matrix-associated proliferation-related antigen. I. Intracellular localization during interphase. J Cell Sci 92 (Pt 1):123–30.

6. Kill IR. 1996. Localisation of the Ki-67 antigen within the nucleolus. Evidence for a fibrillarin-deficient region of the dense fibrillar component. J Cell Sci 109 (Pt 6):1253–1263.

7. Cheutin T, O'Donohue M-F, Beorchia A, Klein C, Kaplan H, Ploton D. 2003. Three dimensional organization of pKi-67: a comparative fluorescence and electron tomography study using FluoroNanogold. J Histochem Cytochem 51:1411–1423.

8. Verheijen R, Kuijpers HJ, van Driel R, Beck JL, van Dierendonck JH, Brakenhoff GJ, Ramaekers FC. 1989. Ki-67 detects a nuclear matrix-associated proliferation-related antigen. II. Localization in mitotic cells and association with chromosomes. J Cell Sci 92 (Pt 4):531–540.

9. Saiwaki T, Kotera I, Sasaki M, Takagi M, Yoneda Y. 2005. In vivo dynamics and kinetics of pKi-67: Transition from a mobile to an immobile form at the onset of anaphase. Exp Cell Res 308:123–134.

10. Takagi M, Nishiyama Y, Taguchi A, Imamoto N. 2014. Ki67 antigen contributes to the timely accumulation of protein phosphatase 1? on anaphase chromosomes. J Biol Chem 289:22877–22887.

11. Booth DG, Takagi M, Sanchez-Pulido L, Petfalski E, Vargiu G, Samejima K, Imamoto N, Ponting CP, Tollervey D, Earnshaw WC, Vagnarelli P. 2014. Ki-67 is a PP1-interacting protein that organises the mitotic chromosome periphery. Elife 3:e01641

12. Sobecki M, Mrouj K, Camasses A, Parisis N, Nicolas E, Lleres D, Gerbe F, Prieto S, Krasinska L, David A, Eguren M, Birling MC, Urbach S, Hem S, Dejardin J, Malumbres M, Jay P, Dulic V, Lafontaine DLJ, Feil R, Fisher D. 2016. The cell proliferation antigen Ki-67 organises heterochromatin. Elife 5:e13722.

13. Van Hooser AA, Yuh P, Heald R. 2005. The perichromosomal layer. Chromosoma 114:377–388.

14. Booth DG, Beckett AJ, Molina O, Samejima I, Masumoto H, Kouprina N, Larionov V, Prior IA, Earnshaw WC. 2016. 3D-CLEM Reveals that a Major Portion of Mitotic Chromosomes Is Not Chromatin. Mol Cell 64:790–802.

15. Cuylen S, Blaukopf C, Politi AZ, Müller-Reichert T, Neumann B, Poser I, Ellenberg J, Hyman AA, Gerlich DW. 2016. Ki-67 acts as a biological surfactant to disperse mitotic chromosomes. Nature 535:308–312.

16. Kumar GS, Gokhan E, De Munter S, Bollen M, Vagnarelli P, Peti W, Page R. 2016. The Ki-67 and RepoMan mitotic phosphatases assemble via an identical, yet novel mechanism. Elife 5:e16539.

17. Schluter C, Duchrow M, Wohlenberg C, Becker MHG, Key G, Flad - HD, Gerdes J. 1993. The cell proliferation-associated antigen of antibody Ki-67: A very large, ubiquitous nuclear protein with numerous repeated elements, representing a new kind of cell cycle-maintaining proteins. J Cell Biol 123:513–522.

18. Kausch I, Lingnau A, Endl E, Sellmann K, Deinert I, Ratliff TL, Jocham D, Sczakiel G, Gerdes J, Böhle A. 2003. Antisense treatment against Ki-67 mRNA inhibits proliferation and tumor growth in vitro and in vivo. Int J Cancer 105:710–716.

19. Zheng J, Ma T, Cao J, Sun X, Chen J, Li W, Wen R, Sun Y, Pei D. 2006. Knockdown of Ki-67 by small interfering RNA leads to inhibition of proliferation and induction of apoptosis in human renal carcinoma cells. Life Sci 78:724–9.

20. Cidado J, Wong HY, Rosen DM, Cimino-mathews A, Garay JP, Fessler AG, Rasheed ZA, Hicks J, Cochran RL, Croessmann S, Zabransky DJ, Mohseni M, Beaver JA, Chu D, Cravero K, Christenson ES, Medford A, Mattox A, De Marzo AM, Argani P, Chawla A, Hurley PJ, Lauring J, Park BH. 2016. Ki-67 is required for maintenance of cancer stem cells but not cell proliferation. Oncotarget 7:6281–6293.

21. Smith CL, Matheson TD, Trombly DJ, Sun X, Campeau E, Han X, Yates JR, Kaufman PD. 2014. A separable domain of the p150 subunit of human chromatin assembly factor-1 promotes protein and chromosome associations with nucleoli. Mol Biol Cell 25:2866–81.

22. Matheson TD, Kaufman PD. 2017. The p150N domain of Chromatin Assembly Factor-1 regulates Ki-67 accumulation on the mitotic perichromosomal layer. Mol Biol Cell 28 (1):21–29.

23. Bodnar AG, Ouellette M, Frolkis M, Holt SE, Chiu CP, Morin GB, Harley CB, Shay JW, Lichtsteiner S, Wright WE. 1998. Extension of life-span by introduction of telomerase into normal human cells. Science 279:349–52.

24. Zhang LF, Huynh KD, Lee JT. 2007. Perinucleolar Targeting of the Inactive X during S Phase: Evidence for a Role in the Maintenance of Silencing. Cell 129:693–706.

25. Dimitrova DS, Berezney R. 2002. The spatio-temporal organization of DNA replication sites is identical in primary, immortalized and transformed mammalian cells. J Cell Sci 115:4037–4051.

26. Fischer M, Grossmann P, Padi M, DeCaprio JA. 2016. Integration of TP53, DREAM, MMB-FOXM1 and RB-E2F target gene analyses identifies cell cycle gene regulatory networks. Nucleic Acids Res 44:6070–6086.

27. Sadasivam S, DeCaprio JA. 2013. The DREAM complex: master coordinator of cell cycle-dependent gene expression. Nat Rev Cancer 13:585–595.

28. Bertoli C, Skotheim JM, de Bruin RAM. 2013. Control of cell cycle transcription during G1 and S phases. Nat Rev Mol Cell Biol 14:518–28.

29. Manning AL, Yazinski SA, Nicolay B, Bryll A, Zou L, Dyson NJ. 2014. Suppression of genome instability in prb-deficient cells by enhancement of chromosome cohesion. Mol Cell 53:993–1004.

30. Litovchick L, Sadasivam S, Florens L, Zhu X, Swanson SK, Velmurugan S, Chen R, Washburn MP, Liu XS, DeCaprio JA. 2007. Evolutionarily Conserved Multisubunit RBL2/p130 and E2F4 Protein Complex Represses Human Cell Cycle-Dependent Genes in Quiescence. Mol Cell 26:539–551.

31. Schmit F, Korenjak M, Mannefeld M, Schmitt K, Franke C, Von Eyss B, Gagrica S, Hänel F, Brehm A, Gaubatz S. 2007. LINC, a human complex that is related to pRB-containing complexes in invertebrates regulates the expression of G2/M genes. Cell Cycle 6:1903–1913.

32. Xiong Y, Hannon GJ, Zhang H, Casso D, Kobayashi R, Beach D. 1993. p21 is a universal inhibitor of cyclin kinases. Nature 363:210–211.

33. Gartel AL, Radhakrishnan SK. 2005. Lost in transcription: p21 repression, mechanisms, and consequences. Cancer Res 65:3980–3985.

34. Labaer J, Garrett MD, Stevenson LF, Slingerland JM, Sandhu C, Chou HS, Fattaey A, Harlow E. 1997. New functional activities for the p21 family of CDK inhibitors. Genes Dev 11:847–862.

35. Waga S, Stillman B. 1998. Cyclin-dependent kinase inhibitor p21 modulates the DNA primer-template recognition complex. Mol Cell Biol 18:4177–87.

36. El-Deiry WS, Tokino T, Velculescu VE, Levy DB, Parsons R, Trent JM, Lin D, Mercer WE, Kinzler KW, Vogelstein B. 1993. WAF1, a potential mediator of p53 tumor suppression. Cell 75:817–825.

37. Deng C, Zhang P, Harper JW, Elledge SJ, Leder P. 1995. Mice lackingp21CIP1/WAF1 undergo normal development, but are defective in G1 checkpoint control. Cell 82:675–684.

38. Junk DJ, Vrba L, Watts GS, Oshiro MM, Martinez JD, Futscher BW. 2008. Different mutant/wild-type p53 combinations cause a spectrum of increased invasive potential in nonmalignant immortalized human mammary epithelial cells. Neoplasia 10:450–61.

39. Petersen G.B, Therelsen A. J. 1962. Number of nucleoli in female and male human cells in tissue culture. Exp Cell Res 28:590–2.

40. Barr ML, Bertram EG. 1949. A morphological distinction between neurones of the male and female, and the behaviour of the nucleolar satellite during accelerated nucleoprotein synthesis. Nature 163:676.

41. Bourgeois CA, Laquerriere F, Hemon D, Hubert J, Bouteille M. 1985. New data on the in situ position of the inactive X chromosome in the interphase nucleus of human fibroblasts. Hum Genet 69:122–129.

42. Lucchesi JC, Kelly WG, Panning B. 2005. Chromatin Remodeling in Dosage Compensation. Annu Rev Genet 39:615–651.

43. Kohlmaier A, Savarese F, Lachner M, Martens J, Jenuwein T, Wutz A. 2004. A chromosomal memory triggered by Xist regulates histone methylation in X inactivation. PLoS Biol 2:E171.

44. Cao R, Wang L, Wang H, Xia L, Erdjument-Bromage H, Tempst P, Jones RS, Zhang Y. 2002. Role of histone H3 lysine 27 methylation in Polycomb-group silencing. Science 298:1039–1043.

45. Di Croce L, Helin K. 2013. Transcriptional regulation by Polycomb group proteins. Natrure strucutral Mol Biol 20:1147–55.

46. Simon JA, Kingston RE. 2013. Occupying chromatin: Polycomb mechanisms for getting to genomic targets, stopping transcriptional traffic, and staying put. Mol Cell 49:808–24.

47. Beck DB, Burton A, Oda H, Ziegler-Birling C, Torres-Padilla ME, Reinberg D. 2012. The role of PR-Set7 in replication licensing depends on Suv4-20h. Genes Dev 26:2580–2589.

48. Chaligné R, Popova T, Mendoza-Parra MA, Saleem MAM, Gentien D, Ban K, Piolot T, Leroy O, Mariani O, Gronemeyer H, Vincent-Salomon A, Stern MH, Heard E. 2015. The inactive X chromosome is epigenetically unstable and transcriptionally labile in breast cancer. Genome Res 25:488–503.

49. Politz JCR, Scalzo D, Groudine M. 2016. The redundancy of the mammalian heterochromatic compartment. Curr Opin Genet Dev 37:1–8.

50. Kind J, Pagie L, Ortabozkoyun H, Boyle S, De Vries SS, Janssen H, Amendola M, Nolen LD, Bickmore WA, Van Steensel B. 2013. Single-cell dynamics of genome-nuclear lamina interactions. Cell 153:178–192.

51. Koningsbruggen S van, Gierlinski M, Schofield P, Martin D, Barton GJ, Ariyurek Y, Dunnen JT den, Lamond AI. 2010. High-Resolution Whole-Genome Sequencing Reveals That Specific Chromatin Domains from Most Human Chromosomes Associate with Nucleoli. Mol Biol Cell 21:3735–3748.

52. Ragoczy T, Telling A, Scalzo D, Kooperberg C, Groudine M. 2014. Functional redundancy in the nuclear compartmentalization of the late-replicating genome. Nucleus 5:626–35.

53. Ma H, Naseri A, Reyes-Gutierrez P, Wolfe SA, Zhang S, Pederson T. 2015. Multicolor CRISPR lableing of chromosomal loci in human cells. Proc Natl Acad Sci U S A 112:3002–7.

54. Benson EK, Mungamuri SK, Attie O, Kracikova M, Sachidanandam R, Manfredi JJ, Aaronson S a. 2014. p53-dependent gene repression through p21 is mediated by recruitment of E2F4 repression complexes. Oncogene 33:3959–69.

55. Sobecki M, Mrouj K, Colinge J, Gerbe F, Jay P, Krasinska L, Dulic V, Fisher D. 2017. Cell cycle regulation accounts for variability in Ki-67 expression levels. Cancer Res canres.0707.2016.

56. Brugarolas J, Moberg K, Boyd S, Taya Y, Jacks T, Lees JA. 1999. Inhibition of cyclin-dependent kinase 2 by p21 is necessary for retinoblastoma protein-mediated G1 arrest after gamma-irradiation. Proc Natl Acad Sci U S A 96:1002–1007.

57. Chubb JR, Boyle S, Perry P, Bickmore WA. 2002. Chromatin motion is constrained by association with nuclear compartments in human cells. Curr Biol 12:439–445.

58. Dillinger S, Straub T, Nemeth A. 2016. Nucleolus association of chromosomal domains is largely maintained in cellular senescence despite massive nuclear reorganisation. BioRxiv DOI: http://dx.doi.org/10.1101/054908

59. Németh A, Conesa A, Santoyo-Lopez J, Medina I, Montaner D, Péterfia B, Solovei I, Cremer T, Dopazo J, Längst G. 2010. Initial genomics of the human nucleolus. PLoS Genet 6:e1000889.

60. Yang F, Deng X, Ma W, Berletch JB, Rabaia N, Wei G, Moore JM, Filippova GN, Xu J, Liu Y, Noble WS, Shendure J, Disteche CM. 2015. The lncRNA Firre anchors the inactive X chromosome to the nucleolus by binding CTCF and maintains H3K27me3methylation. Genome Biol 16:52.

61. Csankovszki G, Nagy A, Jaenisch R. 2001. Synergism of Xist RNA, DNA methylation, and histone hypoacetylation in maintaining X chromosome inactivation. J Cell Biol 153:773–783.

62. Chun-Kan Chen, Mario Blanco, Constanza Jackson, Erik Aznauryan, Noah Ollikainen, Christine Surka, Amy Chow, Patrick McDonel, Andrea Cerase, Mitchell Guttman. 2016. Xist recruits the X chromosome to the nuclear lamina to enable chromosome-wide silencing. Science 354: 468–472.

63. Benanti JA, Galloway DA. 2004. Normal human fibroblasts are resistant to RAS-induced senescence. Mol Cell Biol 24:2842–52.

64. Cong L, Ran FA, Cox D, Lin S, Barretto R, Habib N, Hsu PD, Wu X, Jiang W, Marraffini LA, Zhang F. 2013. Multiplex genome engineering using CRISPR/Cas systems. Science 339: 819–23.

65. Ran FA, Hsu PD, Wright J, Agarwala V, Scott DA, Zhang F. 2013. Genome engineering using the CRISPR-Cas9 system. Nat Protoc 8:2281–2308.

66. Brinkman EK, Chen T, Amendola M, Van Steensel B. 2014. Easy quantitative assessment of genome editing by sequence trace decomposition. Nucleic Acids Res 42s: e168.

67. Yang D, Buchholz F, Huang Z, Goga A, Chen C-Y, Brodsky FM, Bishop JM. 2002. Short RNA duplexes produced by hydrolysis with Escherichia coli RNase III mediate effective RNA interference in mammalian cells. Proc Natl Acad Sci U S A 99:9942–9947.

68. Zhu H. 2012. Cell Proliferation Assay by Flow Cytometry. Bio-Protocol Bio101:e198.

69. Untergasser A, Cutcutache I, Koressaar T, Ye J, Faircloth BC, Remm M, Rozen SG. 2012. Primer3-new capabilities and interfaces. Nucleic Acids Res 40:e115.

70. Trapnell C, Pachter L, Salzberg SL. 2009. TopHat: Discovering splice junctions with RNA-Seq. Bioinformatics 25:1105–1111.

71. Kim D, Pertea G, Trapnell C, Pimentel H, Kelley R, Salzberg SL. 2013. TopHat2: accurate alignment of transcriptomes in the presence of insertions, deletions and gene fusions. Genome Biol 14:R36.

72. Trapnell C, Williams B a, Pertea G, Mortazavi A, Kwan G, van Baren MJ, Salzberg SL, Wold BJ, Pachter L. 2010. Transcript assembly and abundance estimation from RNA-Seq reveals thousands of new transcripts and switching among isoforms. Nat Biotechnol 28:511–515.

73. Zhu LJ, Gazin C, Lawson ND, Pagès H, Lin SM, Lapointe DS, Green MR. 2010. ChIPpeakAnno: a Bioconductor package to annotate ChIP-seq and ChIP-chip data. BMC Bioinformatics 11:237.

74. Zhu LJ. 2013. Integrative Analysis of ChIP-Chip and ChIP-Seq Dataset. Methods Mol Biol 1067:105–124.

75. Wright SP. 1992. Adjustd p-values for simultaneous inference. Biometrics 48:1005–1013.

76. Passerini V, Ozeri-Galai E, de Pagter MS, Donnelly N, Schmalbrock S, Kloosterman WP, Kerem B, Storchová Z. 2016. The presence of extra chromosomes leads to genomic instability. Nat Commun 7:10754.

## Referencs for Supplemental material

1. Trapnell C, Williams B a, Pertea G, Mortazavi A, Kwan G, van Baren MJ, Salzberg SL, Wold BJ, Pachter L. 2010. Transcript assembly and abundance estimation from RNA-Seq reveals thousands of new transcripts and switching among isoforms. Nat Biotechnol 28:511–515.

2. Fischer M, Grossmann P, Padi M, DeCaprio JA. 2016. Integration of TP53, DREAM, MMB-FOXM1 and RB-E2F target gene analyses identifies cell cycle gene regulatory networks. Nucleic Acids Res 44:6070–6086.

3. Wright SP. 1992. Adjustd p-values for simultaneous inference. Biometrics 48:1005–1013.

4. Passerini V, Ozeri-Galai E, de Pagter MS, Donnelly N, Schmalbrock S, Kloosterman WP, Kerem B, Storchová Z. 2016. The presence of extra chromosomes leads to genomic instability. Nat Commun 7:10754.

5. Carrel L, Willard HF. 2005. X-inactivation profile reveals extensive variability in X1298 linked gene expression in females. Nature 434:400–404.

6. Zhang Y, Castillo-Morales A, Jiang M, Zhu Y, Hu L, Urrutia AO, Kong X, Hurst LD. 2013. Genes that escape X-inactivation in humans have high intraspecific variability in expression, are associated with mental impairment but are not slow evolving. Mol Biol Evol 30:2588–2601.

